# Biophysical and Architectural Mechanisms of Subthalamic Theta under Response Conflict

**DOI:** 10.1101/2021.11.10.468101

**Authors:** Prannath Moolchand, Stephanie R. Jones, Michael J. Frank

## Abstract

The cortico-basal ganglia circuit is needed to suppress prepotent actions and to facilitate controlled behavior. Under conditions of response conflict, the frontal cortex and subthalamic nucleus [STN] exhibit increased spiking and theta band power, which are linked to adaptive regulation of behavioral output. The electrophysiological mechanisms underlying these neural signatures of impulse control remain poorly understood. To address this lacuna, we constructed a novel large-scale, biophysically principled model of the subthalamopallidal [STN-Globus Pallidus externus (GPe)] network, and examined the mechanisms that modulate theta power and spiking in response to cortical input. Simulations confirmed that theta power does not emerge from intrinsic network dynamics but is robustly elicited in response to cortical input as burst events representing action selection dynamics. Rhythmic burst events of multiple cortical populations, representing a state of conflict where cortical motor plans vacillate in the theta range, led to prolonged STN theta and increased spiking, consistent with empirical literature. Notably, theta band signaling required NMDA, but not AMPA, currents, which were in turn related to a triphasic STN response characterized by spiking, silence and bursting periods. Finally, theta band resonance was also strongly modulated by architectural connectivity, with maximal theta arising when multiple cortical populations project to individual STN “conflict detector” units, due to an NMDA-dependent supralinear response. Our results provide insights into the biophysical principles and architectural constraints that give rise to STN dynamics during response conflict, and how their disruption can lead to impulsivity and compulsivity.

## Introduction

Various studies have implicated the cortico-subthalamic “hyperdirect-pathway” in response inhibition upon encountering response conflict (Aron et al. 2007; Baunez and Robbins 1997; Baunez et al. 2001; Cavanagh et al. 2011; Frank 2006; Herz et al. 2016; Isoda and Hikosaka 2007, 2008; Kelley et al. 2018; Nambu et al. 2002; Schmidt et al. 2013; Verbruggen and Logan 2008; Wessel et al. 2019; Zaghloul et al. 2012; Zavala et al. 2014). The STN mediates these functions via global suppression of motor output, putatively via diffuse projections to Basal-Ganglia [BG] output structures (Frank 2006; Parent and Hazrati 1995; Wessel et al. 2019). Disruption of STN causes impulsive behavior (Baunez et al. 1995; Baunez and Robbins 1997; Baunez et al. 2001; Coulthard et al. 2012; Fife et al. 2017; Frank et al. 2007; Ghahremani et al. 2018; Green et al. 2013; Wylie et al. 2010). Accounting for these observations, computational models of BG (Bogacz and Gurney 2007; Frank 2006; Ratcliff and Frank 2012; Wiecki and Frank 2013) have further posited that elevated STN firing dynamically raises a “decision-threshold” to afford time to resolve the conflict.

Despite the growing body of work on STN dynamics, the electrophysiological mechanisms remain poorly understood. Many studies have reported elevated theta (4-8Hz) power in frontal-cortex and STN during response conflict (Cavanagh et al. 2011, 2014; Herz et al. 2016; Zavala et al. 2014; Zavala et al. 2013, 2016). Such theta modulations induce transient global suppression of motor activity (Wessel et al. 2019), leading to slower but more accurate choices, as captured by elevated decision-thresholds in quantitative models (Cavanagh et al. 2011; Herz et al. 2017; Herz et al. 2016; Kelley et al. 2018; Zavala et al. 2014; Zavala et al. 2016). But what are the mechanisms of such STN theta-band signals, and how are they modulated? For example, while STN theta correlates positively with decision-threshold and response-time slowing during high conflict conditions, the opposite has been observed during low conflict (Cavanagh et al. 2011; Herz et al. 2016), thereby questioning whether theta power *per se* indicates for the need for response caution. Moreover, many factors could influence STN theta, from conductance dynamics of individual cells (Bevan and Wilson 1999; Cooper and Stanford 2000; Deister et al. 2013; Jones 2016; Rubin 2017; Wilson 2010, 2013) to network topography (Alkemade et al. 2015; Haynes and Haber 2013; Kita et al. 2014; Mathai and Smith 2011; Nambu 2011; Schroll and Hamker 2013), to the dynamics of the cortical inputs to STN (Zavala et al. 2014).

To query the conditions under which STN theta power is modulated, we developed a novel biophysically principled large-scale model of the subthalamopallidal circuit by adapting previous STN and GPe cellular models (Rubin and Terman 2004; Terman et al. 2002). Our model incorporates arkypallidal GPe units, and imposes sparse and probabilistic connectivities (Abdi et al. 2015; Baufreton et al. 2009; Bevan et al. 2007; Kita 2010; Kita 2007; Mallet et al. 2012; Oorschot 1996; Sadek et al. 2007; Wilson 2010). We first constrained our network to capture single cell electrophysiological patterns. We then characterized the effects of various dynamic regimes of cortical inputs (Jones 2016) on STN theta, and how they are altered by cellular mechanisms (e.g. AMPA and NMDA currents). We simulated response conflict in terms of coactivation of multiple cortical populations (representing competing motor responses), and explored the impact of their topographical connectivities to STN subpopulations. Finally, we explored how these mechanisms impact both oscillatory dynamics and spiking in these subpopulations.

Our simulations implicate NMDA-dependent mechanisms within STN in conflict-induced elevations in theta power and spiking. Theta-band resonance was also strongly modulated by architectural constraints, with maximal response achieved when multiple cortical inputs converged on STN subpopulations. Analysis of the underlying mechanism revealed an NMDA-dependent supralinear response in STN “conflict-detector” units. These findings provide a potential resolution to various conundrums in the empirical literature and a mechanistic basis to guide potential therapeutic developments for impulsive behaviors.

## Materials & Methods

### Overview of Subthalamoapallidal [STN-GPe] Network

We constructed a large-scale model of the STN-GPe network, Figure 1A, to study the conditions regulating STN theta and spiking during response conflict. We began by adapting prior models derived from rat electrophysiology of the STN and GPe (Rubin and Terman 2004; Terman et al. 2002), expanding them to include (i) two distinct subpopulations of GPe, (ii) NMDA currents, and (iii) various types of cortical inputs. STN activity is predominantly patterned by reciprocal excitatory and inhibitory interactions with the GPe (Baufreton et al. 2009; Wilson 2010, 2013), and we thus considered it important to incorporate the recently established arkypallidal [GPeA] subclass dynamics and connectivities (Abdi et al. 2015; Mallet et al. 2012), in addition to the prototypic pallidal [GPeP] cells. Because we focus on the subthalamopallidal network, omitting the larger circuit (e.g. striatal input, globus pallidus internus output) in which it is embedded, we began by tuning the baseline state of the network (its intrinsic state with no cortical drives, section Baseline Tuning), to reflect *in vivo* spiking statistics. Whilst capturing the overall effects of the non-modeled nuclei, this trade-off facilitated our model tuning to be challenged by biophysical constraints (see section Biophysical Constraints) of the STN and GPe units at various levels.

**Figure 1:**
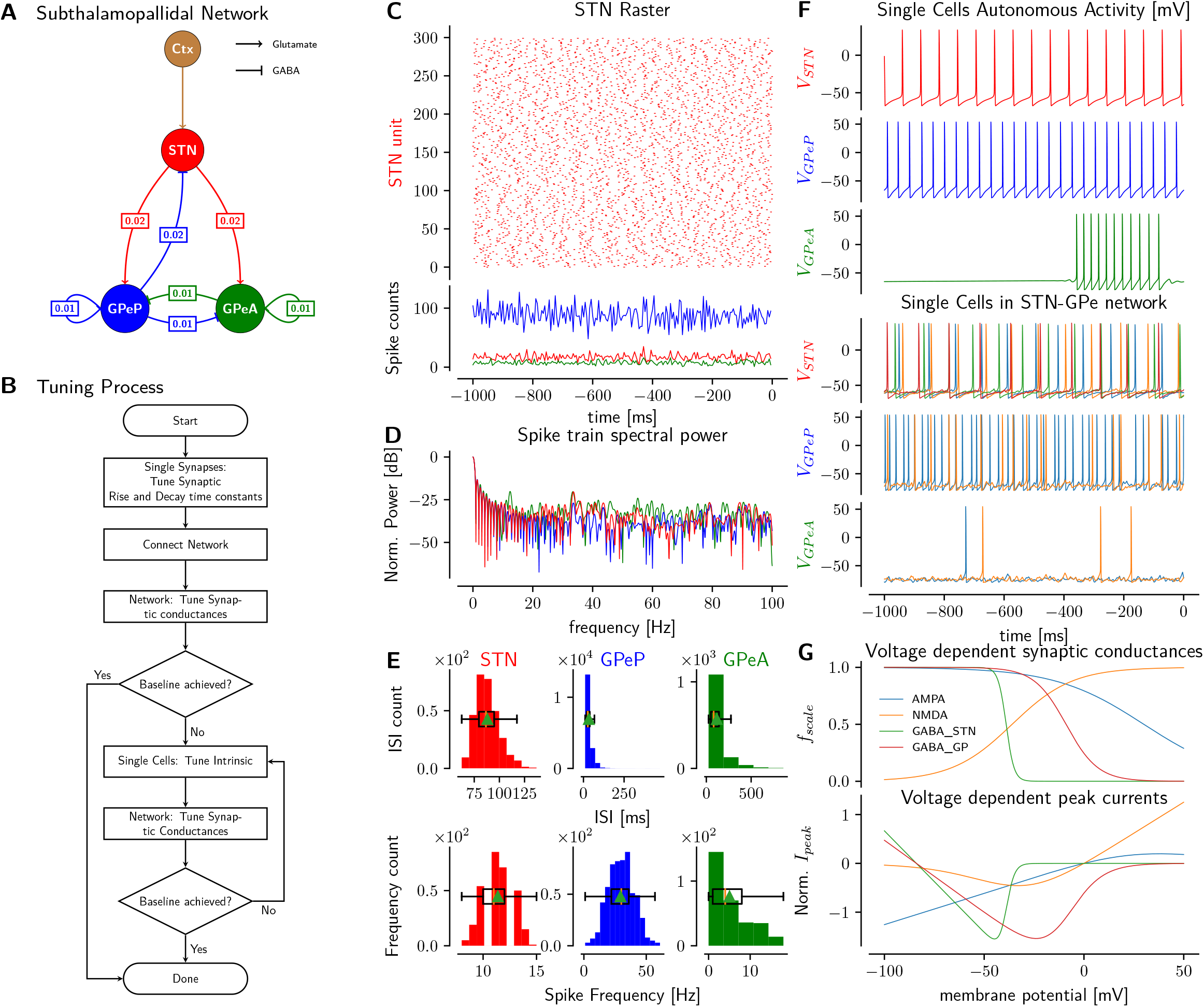
**A:** Subthalamopallidal [STN-GPe] network models with stochastic connectivities; boxed numbers represent connection probabilities. STN - Subthalamic Nucleus, GPeP - Globus Pallidus externus Prototypic, GPeA - Globus Pallidus externus Arkypallidal, Ctx - Cortex [spike trains]. **B:** Tuning process to achieve asynchronous baseline [no cortical drives] network spiking. **C:** Asynchronous baseline network spiking with STN raster and spike counts. Colors as in A, time aligned to cortical drive. **D:** Normalized spectral power of spike trains (in C). **E:** *Upper:* Inter-Spike-Interval and *Lower:* Spiking Frequency counts of subpopulations in baseline network. **F:** *Upper:* Autonomous cellular activities after tuning, in absence of any synaptic inputs. *Lower:* Typical cellular activities in baseline network. **G:** *Upper:* Normalized voltage dependent synaptic conductances after tuning, for AMPA, NMDA and GABA_STN currents on STN, and GABA_GP on GPeP and GPeA. *Lower:* Corresponding current-voltage [I-V] characteristics, normalized to some peak currents.

**Figure 2:**
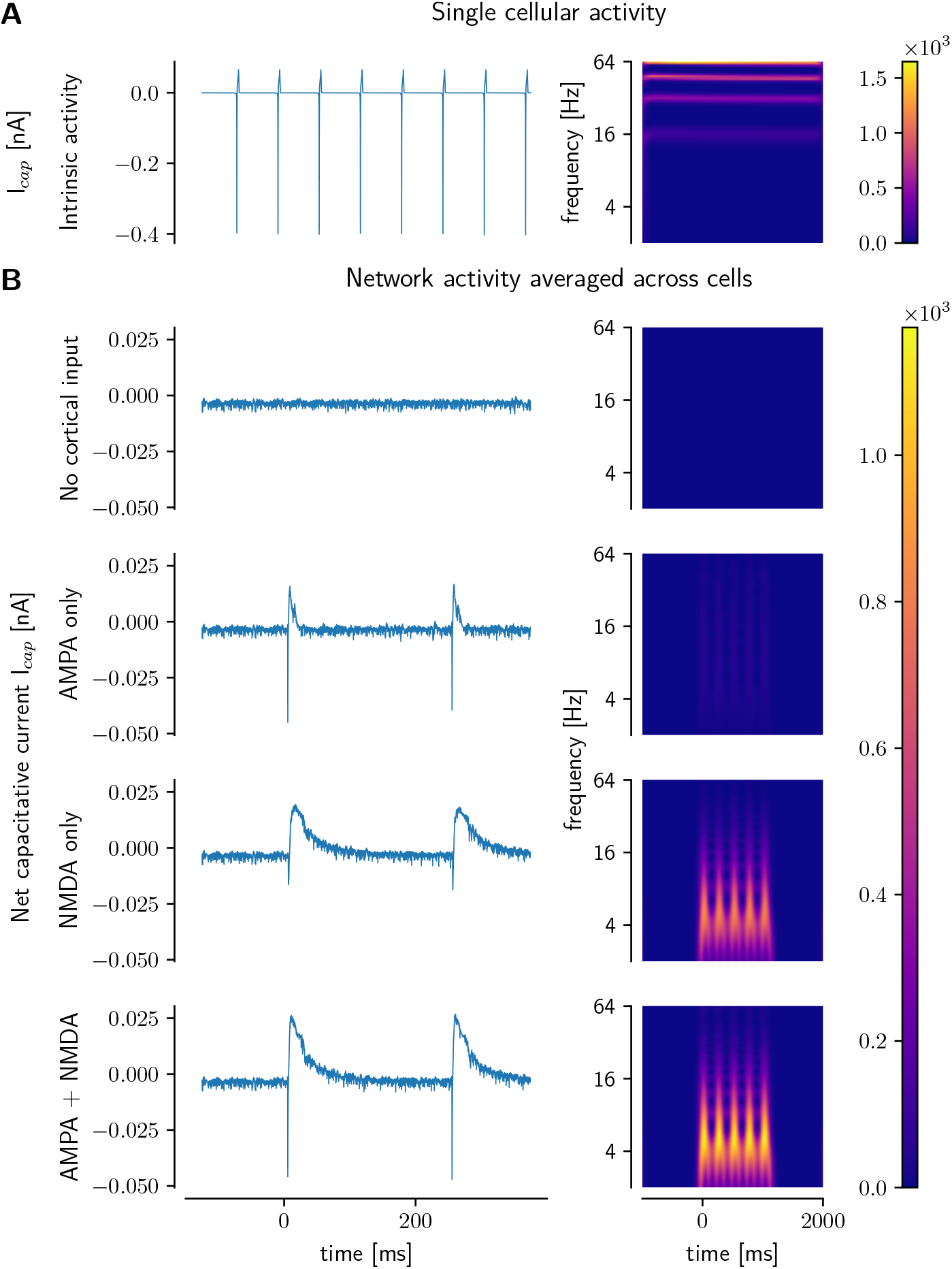
**A:** Celluar activity of single unit after tuning. *Left:* Capacitative current (no synaptic components). *Right:* Spectrogram of capacitative current. **B:** Network activity by averaging capacitative currents across all STN units, under cortical inputs and *in silico* pharmacological blockade, aligned to cortical input onset.

Our simulations were implemented using the Neuron framework in Python with Message Passing Interface Parallelization and Parallel Context (Hines et al. 2009; Jones et al. 2009), and were run on High Performance Computing resources at Brown University and XSEDE resources (Carnevale et al. 2014; Sivagnanam et al. 2013; Towns et al. 2014). Data were stored in the HDF5 (Folk et al. 2011) hierarchical strcuture with Parallel Input/Output functionality.

### Biophysical constraints and network size

Spanning various levels of neuroscience, this study demanded that mechanisms regarding synaptic kinetics, cellular dynamics, intra- and inter-nuclei connectivities conform to biophysical constraints before investigating how these mechanisms may give rise to neural signatures of response conflict. Given the inherent complex nonlinearity across and within several scales, and the compounding tight coupling of biophysical objectives, tuning such a network necessitated several iterative steps, Figure 1B, as described in section Tuning Overview.

### Electrophysiological constraints

We first imposed that postsynaptic currents reproduce the rise and decay kinetics reported under voltage clamp experiments, and that synaptic conductances show voltage dependence, Figure 1G. At the cellular level, we ensured that membrane potentials exhibit physiologically plausible dynamics related to action potential [AP] generation and resting membrane potentials [RMP]; those that looked pathological (such as elevated RMP, APs with broad spike widths and hyperpolarizing peak voltages) were rejected. We also insisted that each subclass of cells accorded with *in vivo* population spiking statistics in terms of means and distribution shapes, Figure 1E, subject to connectivity constraints. Furthermore, because synchronous STN and GPe firing is characteristic of pathological conditions (Eusebio et al. 2009; Goldberg 2004; Hamani 2004; Hammond et al. 2007; Schwab et al. 2013; Wilson 2010, 2013), minimizing the incidence of such synchrony was an additional biophysical constraint on the target baseline state, which has also been captured by other computational models (Hahn and McIntyre 2010; Park et al. 2011; Pavlides et al. 2015). These requirements, coupled with the need to have physiologically relevant temporal resolution to model theta signals, motivated our use of active conductance Hodgkin-Huxley based models (Rubin and Terman 2004; Terman et al. 2002) for somata, see section Cellular Tuning. Because available data from paired recordings were measured at somatic levels only, with no information as regards dendritic ion channel distributions and post-synaptic-potential attenuations, we chose single compartment models.

### Anatomical constraints

The sparsity in subthalamopallidal connectivities (Baufreton et al. 2009; Wilson 2010), Figure 1A, required a large network size. However, the stiff dynamics of the GPe cells dictated a very small simulation step size of 0.0025 ms for numerical accuracy, which placed an upper limit on the network size given our computational resources. These conflicting restrictions, in conjunction with reported anatomical population ratios of STN to GPe being one to three across several species (Hardman et al. 2002; Oorschot 1996), informed our network size of 300 STN and 900 GPe cells. Additionally, prototypic GPe [GPeP] cells are estimated to be around twice the population size of arkypallidal [GPeA] cells (Abdi et al. 2015; Mallet et al. 2012). We thus divided the GPe population into 600 GPeP and 300 GPeA units.

### Overview of the tuning process

Figure 1B provides a flow-chart of the model tuning process. We started by tuning the synaptic kinetics to reproduce the rise and decay of post-synaptic currents under voltage-clamp experiments recorded at the somata, see section Synaptic Tuning. Using the GPe model in Rubin and Terman (2004) as a template, that would correspond to the prototypic subclass in our case, we changed the intrinsic properties to derive the arkypallidal [GPeA] model. Fixing the connection probabilities of the STN and GPeP (Baufreton et al. 2009; Wilson 2010) and our new GPeA, as in Figure 1A, we then performed a thorough search in the synaptic conductance space, and evaluated whether the baseline states of the generated networks met the imposed *in vivo* spiking statistics, Figure 1E. Unfortunately, we found no parameter sets from the original Rubin and Terman (2004) model that could conform to the latter constraints. Consequently, we tuned the dynamics of all the cellular units, namely STN, GPeP and GPeA (i.e. departing from those used by Rubin and Terman (2004)), and repeated the synaptic conductance space search and evaluations. The latter paired tuning process of cellular dynamics followed by synaptic conductance was iterated until a parameter set that met several biophysical constraints was obtained. We note here that several parameter sets, while successful in capturing the desired spiking statistics, were eventually rejected when challenged by biophysical constraints at the cellular level, such as action potential characteristics (see section Cellular Tuning).

### Tuning synaptic dynamics

Lack of data to tune metabotropic postsynaptic influences on single cells restricted our investigations to ionotropic currents, Equation 1a, only.

In contrast to solving the first-order differential model for synaptic activity in Rubin and Terman (2004) and Terman et al. (2002), we opted for the computationally less expensive biexponential model with rise and decay constants, Neuron’s Exp2Syn mechanism (Carnevale and Hines 2006), Equation 1b, for synaptic responses, which we modified to allow maximal conductances 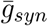 to scale with voltage, Equation 1e, to capture more realistic current-voltage [I-V] characteristics (Chu et al. 2015). This **novel formalism** compensates for the diminishing depolarizing driving force *E_glutamatergic_* — *V* of synaptic currents as the cell depolarizes, which is particularly influential for NMDA dynamics and also exhibits the inward rectification inherent in cortico-STN AMPAergic activities (Chu et al. 2015), see Figure 1G. To control the dynamic range, saturation and positive/negative relationship for the voltage-dependent scaling of conductance, we adopted a logistic model, Equation 1e, with parameters described in Table 1. Despite the continuous dependence of the maximal conductance 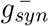 on *V*, the current is non-zero only when triggered by Neuron’s NetCon mechanism until it decays based on its kinetics; this triggering is activated when the presynaptic unit crosses the specified spike threshold and sends a message to the NetCon to start solving the differential equations for the synaptic currents (Carnevale and Hines 2006). Relatedly, we also specified the synaptic transmission delays *δ_syn_* in the NetCon mechanism. Moreover, since we discarded parameter sets with unrealistic membrane potentials, the scaling factor and hence the synaptic conductances remained within physiological bounds.

**Table 1:**
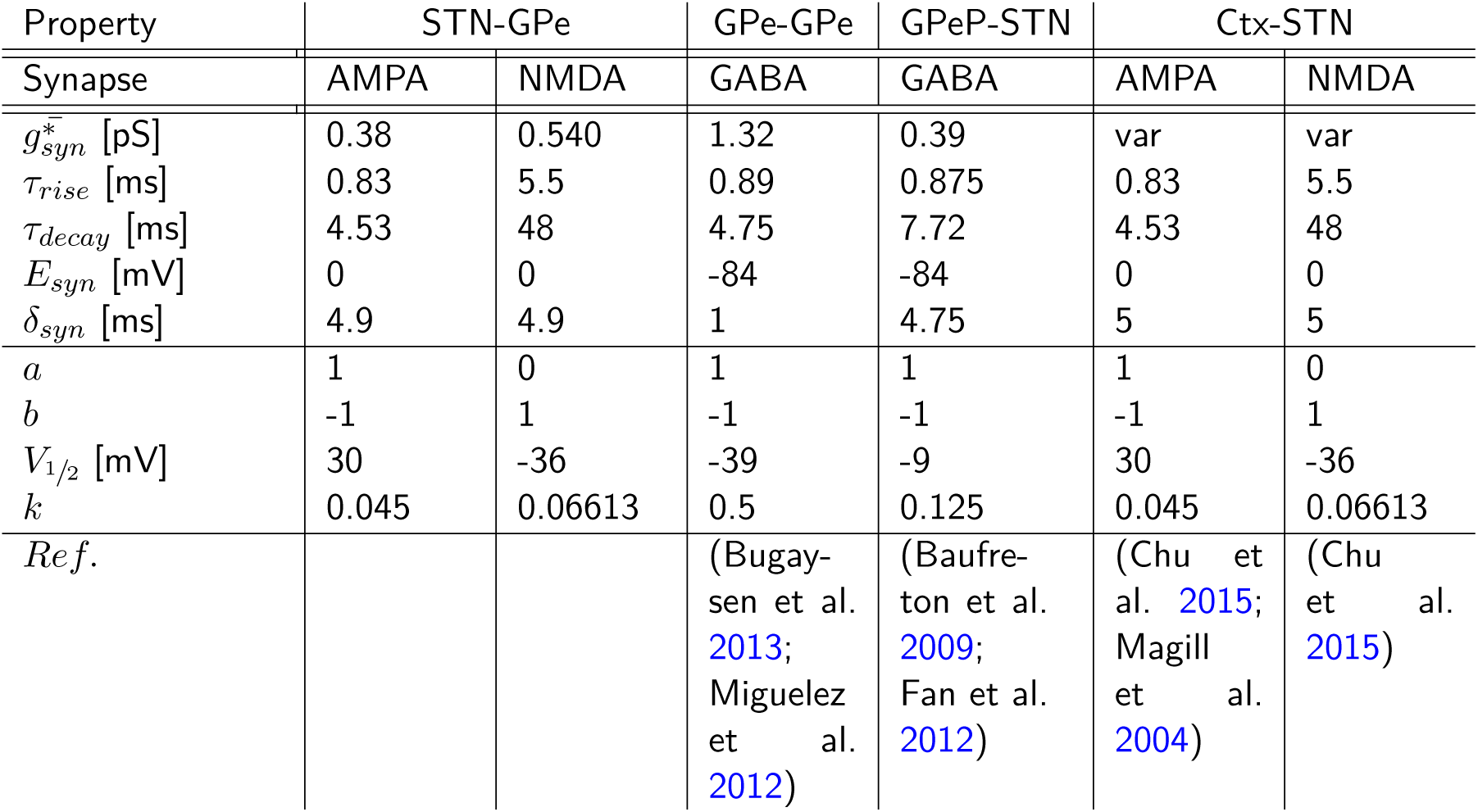
Parameters for synaptic models. Key: GPe parameters apply to both GPeP and GPeA subpopulations; GPeP refers to the prototypic GPe only. Due to unavailability of data, we have assumed the kinetics and voltage-dependent conductance for the glutamatergic synapses on GPe subpopulations to be the same as those on the STN.

#### Postsynaptic current

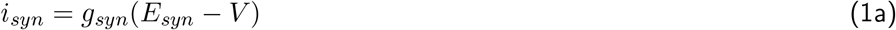

where *i, g* and *E* are the current, conductance and reversal potential respectively

#### Closed-form solution for bi-exponential model of synaptic conductance

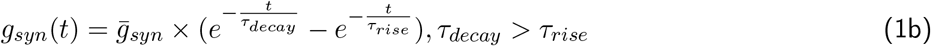

where *g* and 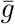 are the instantaneous and maximal synaptic conductances, respectively

#### Voltage dependent maximal synaptic conductance model

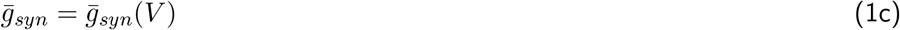

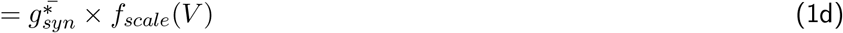

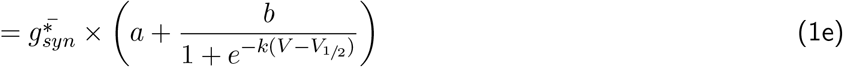

where 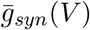 and 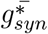 are the voltage-dependent maximal synaptic conductance and reference conductance, respectively. For different synapses, 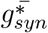’*s* are either tuned to achieve baseline conditions (see section Baseline Tuning) or are manipulated to investigate cortico-STN interactions. *f_scale_*(*V*) is the voltage-dependent scaling factor, modeled by the logistic function.

Despite its simplicity, the biexponential model does not have a closed-form solution to solve for the (10-90%) rise *τ_rise_* and decay *τ_decay_* rates. To replicate the time courses of synaptic responses under voltage clamps, we manually fitted the biexponential model by guessing an initial solution of time rises and decays to achieve the 10-90% characteristics, iteratively refining the ranges to converge upon a satisfying solution, which are reported in Table 1.

### Individual Cell Dynamics and Synaptic Connections

STN and GPeP cells were modeled as in Terman et al. (2002), with the *Na*^+^, *K*^+^, *leak*, *L-type Ca*^2+^, *T-type Ca*^2+^, and [*Ca*^2+^]-*dependent K*^+^ currents and active conductances (Equation 2). Note that our GPeP units would correspond to the GPe described in Terman et al. (2002). We then made parameter adjustments to model the novel GPeA cells to capture their lower firing rates and subpopulation sizes, and also to incorporate the fact that only the GPeP, but not the GPeA, send reciprocal inhibition to the STN, see Figure 1. This latter anatomical difference critically affects STN-GPe theta and spiking dynamics, thus justifying the partition of the GPe into two subclasses.

All equations and parameters for STN and GPeP were as in Rubin and Terman (2004) and Terman et al. (2002), up to specific capacitance *C_m_* scaling to 1 *μF/cm*^2^ for every cell, except for changes detailed in Table 2.

**Table 2:** Parameters for cellular spontaneous activity. Symbols are same as in Terman et al. (2002). The dynamics of the novel GPe Arkypallidal (GPeA) cells have been adapted from the tuned Prototypic (GPeP) cells; only the latter were included in Terman et al. (2002).

| Description | Parameter | Value | Unit |
| --- | --- | --- | --- |
| STN $Na^+$ conductance | $g_{Na}$ | 55 | $mS/cm^2$ |
| STN $K^+$ conductance | $g_K$ | 30 | $mS/cm^2$ |
| STN $Na^+$ $m$-gate half activation voltage | $\theta_m$ | -37 | $mV$ |
| GPeP $Na^+$ conductance | $g_{Na}$ | 120 | $mS/cm^2$ |
| GPeP $K^+$ conductance | $g_K$ | 30 | $mS/cm^2$ |
| GPeP $Na^+$ $h$-gate kinetics factor | $\phi_h$ | 0.05 | |
| GPeP $K^+$ $n$-gate kinetics factor | $\phi_n$ | 0.05 | |
| GPeP $Ca^{2+}$ pump rate | $k_{Ca}$ | 15.0 | |
| GPeA $Na^+$ conductance | $g_{Na}$ | 97 | $mS/cm^2$ |
| GPeA $K^+$ conductance | $g_K$ | 27.5 | $mS/cm^2$ |
| GPeA $Na^+$ $h$-gate kinetics factor | $\phi_h$ | 0.05 | |
| GPeA $K^+$ $n$-gate kinetics factor | $\phi_n$ | 0.05 | |
| GPeA $Ca^{2+}$ pump rate | $k_{Ca}$ | 15.0 | |

#### Membrane dynamics

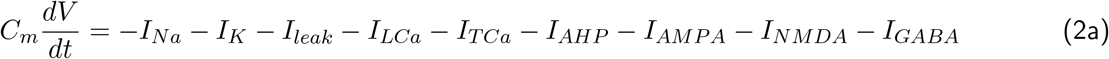

#### Action potential generating currents

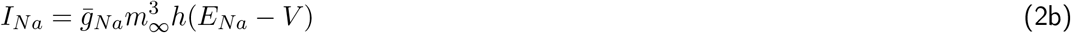

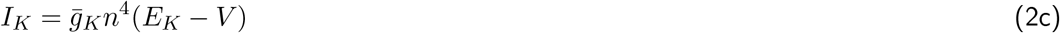

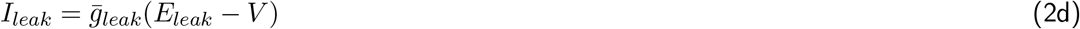

#### *Ca*^2+^ related currents

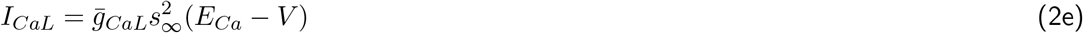

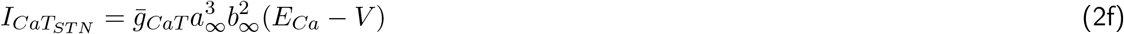

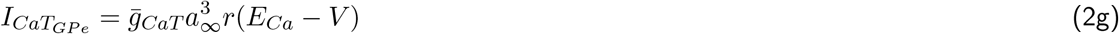

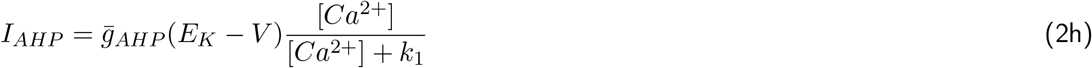

#### Postsynaptic currents

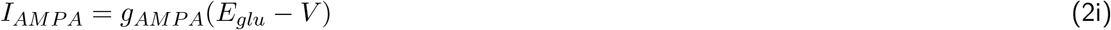

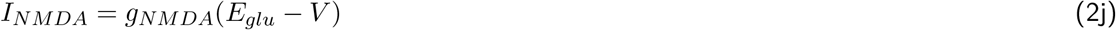

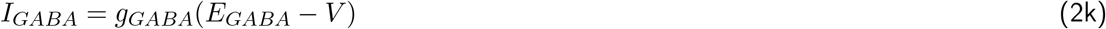

#### First-order kinetics for gating variables: *n,h*

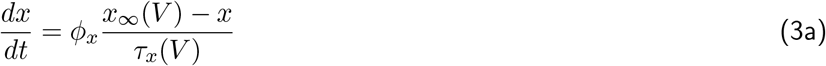

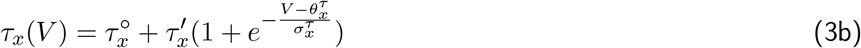

#### Steady state voltage-dependence for gating variables: *m, n, h*

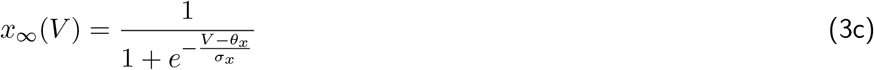

#### [*Ca*^2+^] dynamics

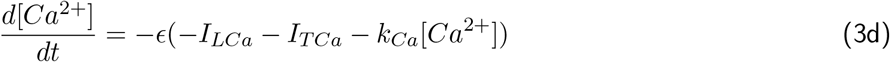

Spike detection thresholds for activating the NetCons were set at −47.4, −56.6 and −55.0 mV for STN, GPeP and GPeA units respectively.

Each STN unit had an AMPA and an NMDA synapses responding to cortical spike trains. The cortico-STN synaptic parameters, Table 1, were matched to data from Chu et al. (2015), both for kinetics and maximal conductance voltage dependence. The NMDA to AMPA maximal conductance ratio was kept at a fixed value corresponding to experimentally measured NMDA conductance at 40 mV divided by the AMPA conductance at −80 mV, which in this case is 1.402. Note that we use *f_scale_* = 1 for NMDA at 40 mV and AMPA at −80 mV in the voltage-dependent maximal conductance formalism, Equation 1e, Figure 1G; hence the maximal conductances would match the experimental values. Each STN unit had a GABA synapse for responding to GPeP stimulation. Both GPeP and GPeA cells had an AMPA and an NMDA that received connections from the STN, and a GABA synapse that was triggered by GPeP and GPeA activities. The different units were probabilistically connected as shown in Figure 1A, with the values in square boxes indicating the sparse connection probabilities, based on previous reports (Abdi et al. 2015; Baufreton et al. 2009; Bevan et al. 2007; Kita 2010; Kita 2007; Mastro et al. 2014; Sadek et al. 2007; Wilson 2010). For every pair of pre- and post-synaptic cells, we performed a Bernoulli trial based on the predefined sparse probabilities, and connected the pair if the result was 1.

### Baseline Tuning of the STN-GPe network

After probabilistically connecting the network, we first tried to reproduce a baseline state that represents a healthy condition, where the units in the network fire asynchronously within and across subpopulations (Figure 1C-F), with normally distributed spiking rates with means for STN, GPeP and GPeA set around 11 (Beurrier et al. 1999; Bevan and Wilson 1999; Kreiss et al. 1997), 30 and 5 spikes/s respectively (Abdi et al. 2015; Benhamou et al. 2012; Cooper and Stanford 2000; Deister et al. 2013; Kita and Kitai 1991; Mallet et al. 2012; Mastro et al. 2014; Nambu and Llinas 1994). The first round of network tuning used the original parameters of the STN model (Terman et al. 2002), which had autonomous firing rate around 2 spikes/s, and could not satisfy the constraints.

Since baseline firing rates depend on both cellular dynamics and synaptic conductances, we first adjusted the intrinsic dynamics of the cells, followed by synaptic parameter searches, iterating these two processes until our biophysical constraints were satisfied (Figure 1B). To further constrain our parameter sets, we analyzed the membrane potentials of the cells in the network and used only those that showed realistic non-pathological voltage traces (Figure 1F), discarding, for instance, those which had any combinations of: very high peak voltages (above 60 mV), hyperpolarized peak spike voltages, exaggerated action-potential half-widths, after-hyperpolarization-potentials lower than −150 mV, resting membrane potentials more depolarized than −50 mV or more hyperpolarized than −85 mV. Final parameters are detailed in Table 2.

### Cortex to STN Stimulation

Our goal in this study was to examine both biophysical and architectural constraints that could give rise to the observed cortico-subthalamic theta power and spike rate modulations during response conflict. To this end, we first investigated cortical patterned activities that support increases in overall STN theta power, followed by how cortico-STN connectivity mediates theta power modulation in the context of conflict.

### Cortico-STN drive

Cortical activity was modeled as spike trains activating STN AMPA and NMDA synapses. However, due to paucity of cortical spiking data in response conflict conditions, we examined two cortical spiking models, Figure 3A, B, that have been shown (Jones 2016) to underlie frequency band power modulation.

**Figure 3:**
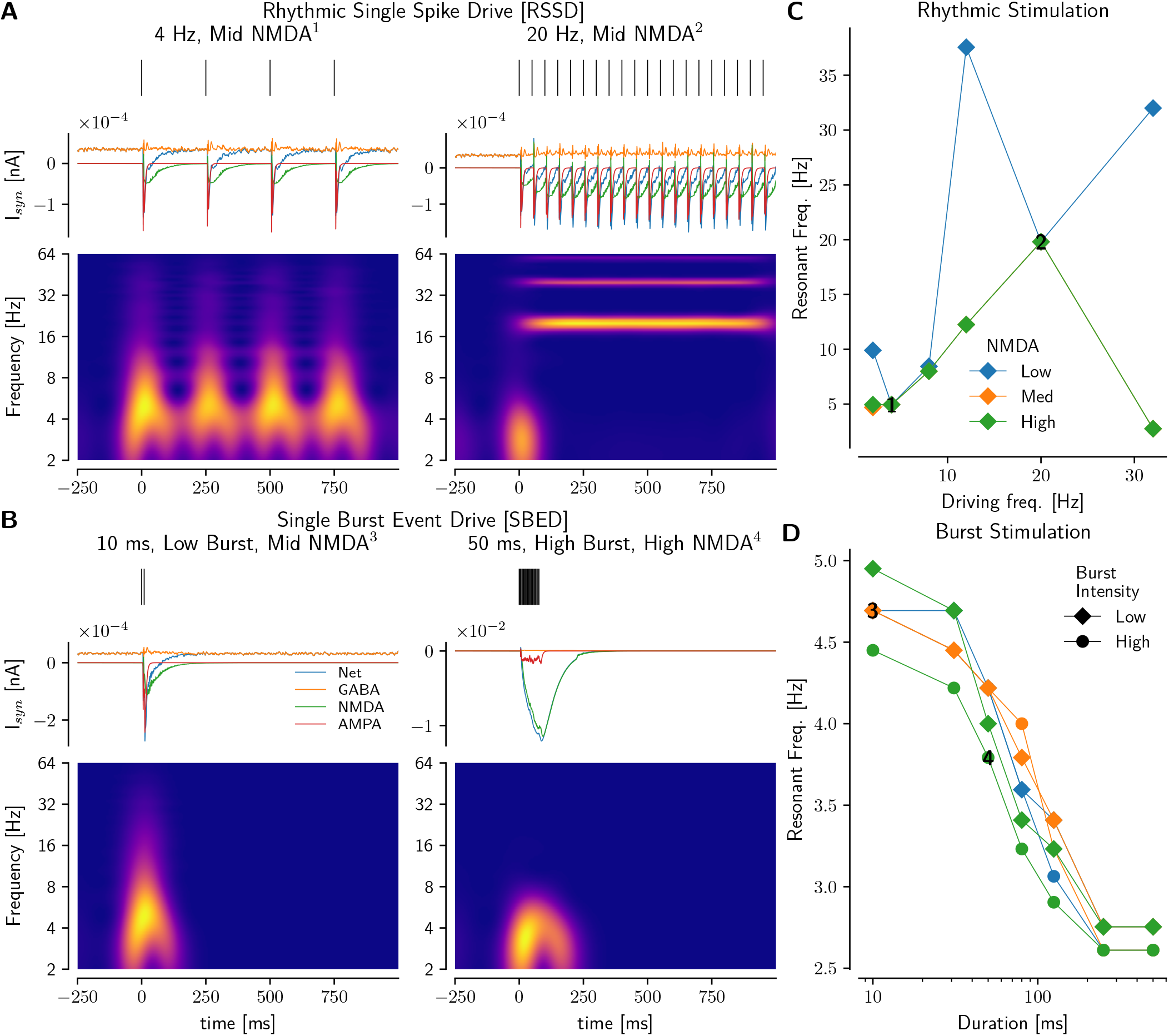
**A:** Rhythmic Single Spike Drive **B:** Single Burst Event Drive. *Top:* Cortical spike train. *Middle:* Synaptic currents (Net: blue, GABA: orange, NMDA: green, AMPA: red, legend in B, left) averaged across STN units in network. *Bottom:* Spectrogram. **C, D:** Resonant frequency (frequency with highest power) for **C:** RSSD as in A, and **D:** SBED stimulations. *Note:* Numbers in C and D show the corresponding cortical drives in A and B. Color intensity scales are different in each spectrogram.

The first cortical spiking profile, Rhythmic-Single-Spike-Drive [RSSD], consisted of single spike trains generated with fixed time periods, *T_per_*, (rhythmic driving frequency, *f_drv_* = 1/*T_per_*), with stimulation time, *T_stim_* ms, Figure 3A (*Left:* [*T_per_* = 250*ms*, *f_drv_* = 4H*z*], *Right*: [*T_per_* = 50*ms*, *f_drv_* = 20H*z*]).

The second spiking class consisted of a burst of spikes over a fixed duration *E_Dur_*, referred to as Single-Burst-Event-Drive [SBED], see Figures 3E and 5B. The inter-spike-intervals [ISI] within the burst were drawn from a Gaussian distribution. Note that in SBED, *E_Dur_* is the same as the stimulation time *T_stim_*. We vary spiking intensity, that is, the number of spikes in the SBED by controlling the means [*IBI_μ_*] and standard deviations [*IBI_σ_*] of the latter Gaussian distributions; thus, (*IBI_μ_, IBI_σ_*) = (9, 6) ms was characterized as Low-Burst-Intensity [*Burst_Low_*], while (*IBI_μ_, IBI_σ_*) = (3, 2) ms constituted High-Burst-Intensity [*Burst_High_*]. For example, Figure 3B *Left* has *E_Dur_* = *T_stim_* = 10 ms, *IBI_μ_* = 9 ms, *IBI_σ_* = 6 ms, characterizing a Low-Burst-Intensity SBED stimulation.

We extended the latter class to a Rhythmic-Burst-Events-Drive [RBED], where SBEDs are generated at fixed time intervals called Inter-Event-Interval Δ, Figure 5B. In this case, the stimulation time, *T_stim_* = *N* × *E_Dur_* + (*N* − 1) × Δ ms, where *N* is the number of SBEDs.

Based on initial observations that replicated physiologically plausible synaptic responses and non-pathological action potentials, we varied the NMDA conductances through two orders of magnitudes, with the values 5.208 × 10^-7^, 5.208 × 10^-6^ and 5.208 × 10^-5^ nS (specific somatic capacitance normalized to 1 *μF/cm*^2^)), referred to as *NMDA_Low_*, *NMDA_Mid_* and *NMDA_High_* respectively.

### Cortico-STN architecture

Because we ultimately modeled response conflict in terms of simultaneous activation of multiple cortical populations representing different motor responses (see section Simulating Conflict), we also considered that these populations could in turn differentially project to STN subpopulations. Indeed, given the somatotopic organization of cortico-STN afferents (Nambu 2011), we divided the STN population into four subpopulations, and implemented two cortical feeds that probabilistically targeted two subpopulations. Since subpopulations conceptually represented somatotopic areas, cortical motor programs coding for mutually exclusive behaviors could thus preferentially innervate their own target STN subpopulations (see Figure 5A). However, the degree to which such coding is completely segregated along cortical-STN “channels” - or whether crosstalk occurs to share cortical information - is unknown.

We thus parametrically varied the connection probability of a cortical feed to its own STN subpopulation. When a given cortical feed representing a single motor response connected exclusively to its own STN subpopulation, consequently segregating the cortico-STN communication, we call it a Segregated Topography, Figure 5, with target probability *p_tar_* = 1 (all the STN units in the subpopulation are homogeneously driven by the same cortical feed) and non-target probability *p_non-tar_* = 0. A Non-Segregated Topography was defined by allowing crosstalk between cortical feeds and STN subpopulations by varying the value of *p_tar_* in the set (0.85, 0.55 and 0.25) under the constraint *p_tar_* + 3 × *p_non-tar_* = 1 to ensure that the total expected synaptic connections stayed the same across all levels of segregation. This constraint disambiguated the contributions of cortico-STN topography from cortical input energy on STN theta power.

Note that *p_tar_* = 1 corresponds to a fully organized columnar topography while *p_tar_* = 0.25 a fully random one. The probabilistic connectivities in the Non-Segregated cases gave rise to STN units that received from both cortical feeds, termed *Conflict Detectors* (Figure 5A *right*), since they could, at the single cell level, listen to conflicting mutually exclusive motor programs. (Note that even in the segregated cases, however, it is still possible for the STN population as a whole to respond to conflict, given that multiple sub-populations would be co-active at once and could also indirectly interact via their reciprocal projections to GPe.)

### Simulating Conflict

To simulate Conflict conditions, we activated both cortical feeds with temporal overlap (Figure 5B). To vary the levels of Conflict, we varied the durations *E_dur_* of the feeds, as well as the delay *δ* and overlap ⋂ between cortical feeds (Figure 5B).

Further, because many experiments report increases in cortical theta itself during response conflict (Cavanagh et al. 2014), which is Granger causal to STN theta (Zavala et al. 2014), we also tested the effects of rhythmic activity in cortical input, termed Rhythmic-Burst-Event-Drives (Figure 5B *right*, section Cortical Drives).

With these methods, we could thus investigate how STN biophysics, cortical-STN architecture, cortical dynamics and their combinations impact STN theta.

### Analysis

Custom Python scripts were written for all analyses and are available upon request.

### Spectral Analysis

Spectral analysis was performed on simulated local field potentials (LFP), which were modeled as net synaptic currents across the population of interest.^1^ Specifically, LFP was calculated as the mean squared of the sums of synaptic currents across all cells in the population of interest at every recorded time point, with units (*nA*)^2^. ^2^

Spectral analysis was performed using the complex Morlet wavelet method, which was generated using Scipy and further normalized to ensure that the net input energy of the Morlet wavelet across various frequencies was controlled for. For every frequency in the range of 2 through 64 Hz (logarithmically distributed), we calculated the complex Morlet signal with 3.5 cycles and 2^17^ (equivalent to around 3.3 s long kernel) points at a sampling period of 0.025 ms to create a time frequency spectrogram.

To calculate theta energy, we integrated the theta power across stimulation time, which we report in units of (*nA*)^2^*ms*.

For efficiency, we opted for spectral domain multiplication using the Scipy *fftconvolve* routine after demeaning our time series data. The amplitudes of the resulting process were squared and treated as spectral power. To compute the power of a specific band, we average over all the frequencies that are members of that band at every time points.

Resonant frequency was defined as the frequency that showed highest power.

### Spike Statistics

All raster plots shown represent the exact recorded spike times (i.e. without binning); the latter were also used to compute the Inter Spike Intervals, ISIs, for every unit in the network.

To compute population spike counts, we first discretized the spiking data of each cell in fixed bin sizes of 5 ms, and aggregates in each bin were taken over all STN units that constituted a population of interest.

For cumulative spiking, we used the above spike counts for each subpopulation and cumulatively add them over succeeding time bins. To estimate the increase in spiking rate over the stimulation period, we divided the total number of spikes over that period by the total duration of the stimulation. Note that for rhythmic bursts, the inter burst intervals were also counted as part of the stimulation duration.

### Statistical Analysis

We performed simple two-tail Welch’s T-test (SciPy Stats) between pairs of conditions, each with matched number of trials, for average theta power, average spiking rates and average theta-spiking, and we used a significance level of 0.05.

## Results

Having tuned our model to reproduce empirical population spike statistics, we set out to explore the impact of synaptic conductances and cortical inputs on LFPs and spikes. We then proceed to model “conflict” in cortical inputs, and how its impact on STN theta may be influenced by different architectural assumptions regarding cortico-STN topography.

### The Subthalamopallidal network requires cortically-driven NMDA currents to show theta band resonance

We first confirmed that theta band activity did not emerge autonomously in the isolated STN unit or the connected network. Indeed, while the autonomous pacemaking activity of the isolated STN unit, i.e. around 16 spikes/s when unconnected, eventually reproduced the empirical average firing rate of 11 spikes/s in conjunction with the inhibitory GPe layer, Figures 1A, C-F, it did not show any theta band resonance. In this setting, the STN unit showed only higher frequency band activities, Figure 2A, mostly reflecting harmonics of approximately 16 Hz activity. To assess whether theta could emerge within the subthalopallidal network (e.g. due to reciprocal GPe interactions; Figure 1A), we averaged the net capacitative currents of all STN units and computed the spectrograms. Once again, no theta band power was observed (Figure 2B). Because the network was probabilistically connected, we ran several (n=15) instantiations; the absence of theta band activity was consistent across all such trials (data not shown; see also Figure 1D).

We next tested if and how cortical inputs to the STN could drive theta band rhythmicity within the network, as suggested empirically (Zavala et al. 2014). Because cortical rhythms are often expressed as transient bursts of activity (Jones 2016), we considered multiple regimes of cortical drive: rhythmic single spike trains or as a burst of activity over a short duration (RSSD and SBED, respectively; see section Cortical Drives and Figure 3 for illustrations).

Because glutamatergic cortical drive could influence STN via both AMPA and NMDA currents, we isolated their contributions by blocking one or the other. These simulations revealed that AMPA currents are insufficient to generate theta band resonance even when cortical inputs are driven at 4 Hz (i.e. in the theta band; Figure 2B). Notably, however, theta resonance emerged with the addition of NMDA synapses in this setting, and persisted with the removal of AMPA currents (Figure 2B). Moreover, these results held also for SBED stimulations. As such, NMDA activity was necessary for theta band resonance in our model. We explore the biophysical mechanisms of this finding below, with particular focus on the NMDA currents, given their slower decay timecourses (i.e. longer relaxation times) that could give rise to potent effects on theta.

### Cortical bursts are filtered by NMDA dynamics to robustly elicit theta band activity

Having established that cortically-driven NMDA currents promote STN theta band activity when stimulated in the theta range, we next examined if theta was a resonant property of the driven network. A standard way to investigate resonance in nonlinear systems with feedback connections is to analyze its response to rhythmic [RSSD] external drives. We drove the network across a range of frequencies (2, 4, 8, 12.5, 20 and 32 Hz). We observed that for mid and high levels of NMDA conductances, the STN showed resonance at the driving frequency (Figure 3C), except at 32 Hz where these higher NMDA currents prevented repolarization and spiking. Interestingly, when the driving frequency was within the theta band (4-8 Hz), the resonant frequency matched the driving frequency for all levels of NMDA conductances. The dynamics of AMPA, NMDA, GABA and net synaptic currents and spectrograms are shown for RSSD examples of 4 and 20 Hz in Figure 3A.

We then tested the effects of cortical burst event patterns (SBED; see section Cortical Drives) as might occur during brief periods of response conflict (Isoda and Hikosaka 2007, 2008) (without yet simulating conflict *per se*; i.e. these simulations were conducted with a single cortical feed). The durations and burst intensities of the events were varied to contrast a ‘Low Stim’ protocol (10 ms duration and inter burst interval of 9 ms, Figure 3B *right*) with a ‘High Stim’ protocol (50 ms duration, 3 ms inter burst interval, Figure 3B *left*). Despite this large variability in burst events, both simulations yielded theta band resonance. Confirming the low pass filtering effect of NMDA currents, the High Stim cases yielded lower resonant frequencues (Figure 3D), due to larger NMDA peak currents’ admitting longer relaxation times.

Notably, while both rhythmic and burst events produced theta band activities, these findings were differentially dependent on NMDA conductances (Figure 3C,D). In the rhythmic driving mode, larger NMDA conductances were needed to support lower frequency resonance (resonance occurred at higher frequencies when NMDA conductances were smaller). In contrast, burst event stimulations yielded theta band resonance over all three levels of NMDA and over a wide range of durations (albeit with lower resonant frequencies with longer event durations and higher burst spiking, as described above). In sum, cortical burst events robustly induced theta band resonance in the network, whereas rhythmic inputs required increasing NMDA conductance to do so.

### Cortically evoked NMDA currents induce both STN burst spiking and silence periods (triphasic response) during theta resonance

What are the spiking characteristics of STN units during theta resonance? Consistent with empirical reports in both primates and rodents (Hamani 2004; Janssen et al. 2017; Kitai and Deniau 1981; Nambu et al. 2000; Pautrat et al. 2018; Schmidt et al. 2013) (see Figure 4E), STN units in our model could respond to cortical stimulation with either burst spiking (Figure 4 *left column*) or with a triphasic spiking response characterized by initial synchronization, followed by a pause (defined as the average of the individual first ISIs that followed the first spiking evoked by cortical stimulation) and then finally bursts (Figure 4 *right column*). The latter pattern was predominant for cortical spike trains that induced longer and larger NMDA currents, which were required for theta resonance. These simulations highlighted a critical role for this silent period in the triphasic response. Indeed, while one might expect that the silent period is induced by feedback inhibition from GPe, it occurred simultaneously with the timecourse of NMDA currents.

**Figure 4:**
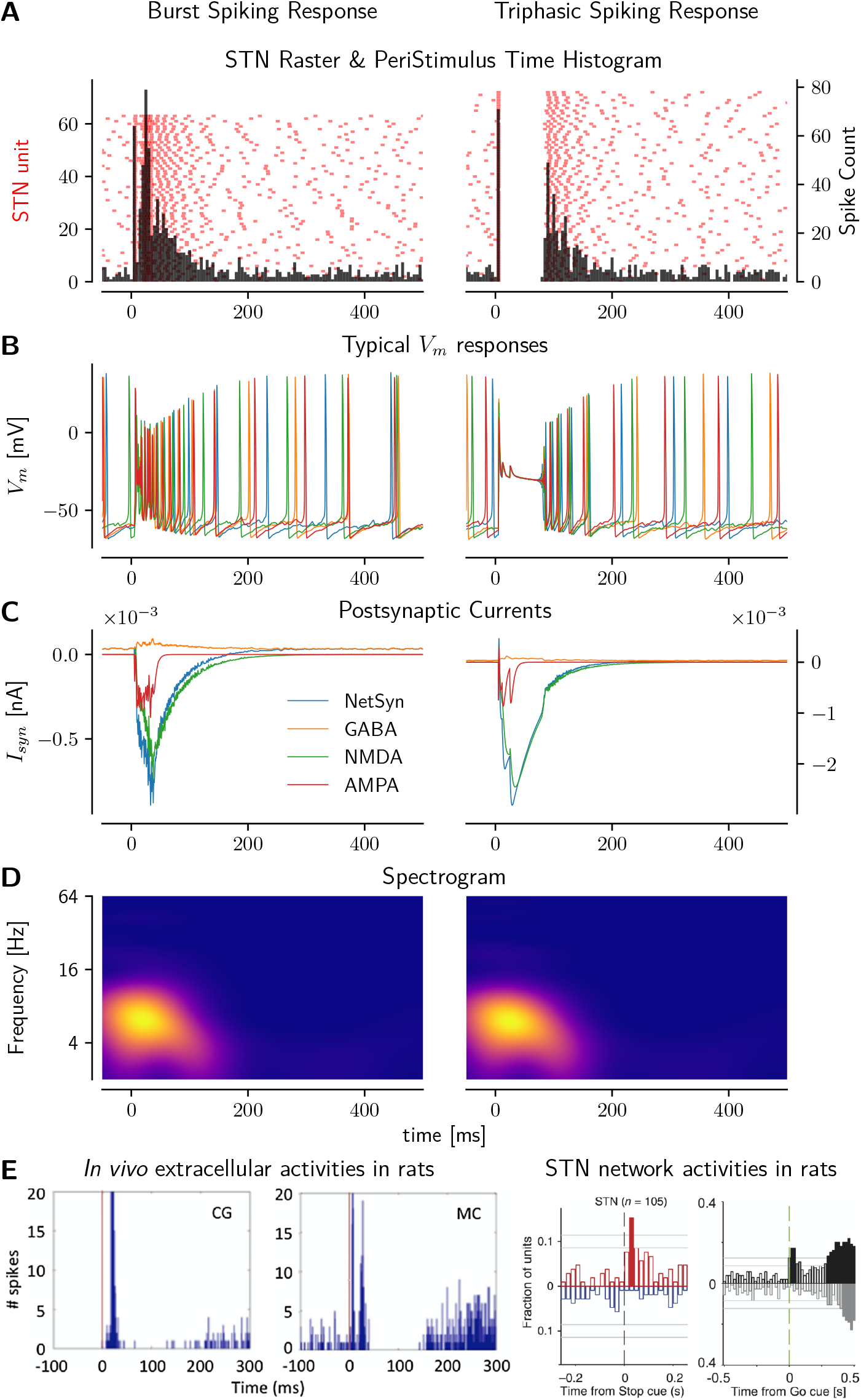
**A-D:** *Left*: Burst Spiking Response. *Right:* Triphasic (Spike Synchronization, Silence, Burst) Spiking Response, in STN subpopulation. **A:** STN raster and PSTH. Left ordinate: STN unit number, right ordinate: Spike count, bin size = 5 ms. **B:** Typical electrophysiological responses of stimulated units. **C:** Postsynaptic currents averaged across all STN units in subpopulation. **D:** Spectrograms (different color intensity scales). **E:** Published *in vivo* activities in rats from extracellular recordings at *Left:* extracellular (Janssen et al. 2017), and *Right:* network (Schmidt et al. 2013) levels.

**Figure 5:**
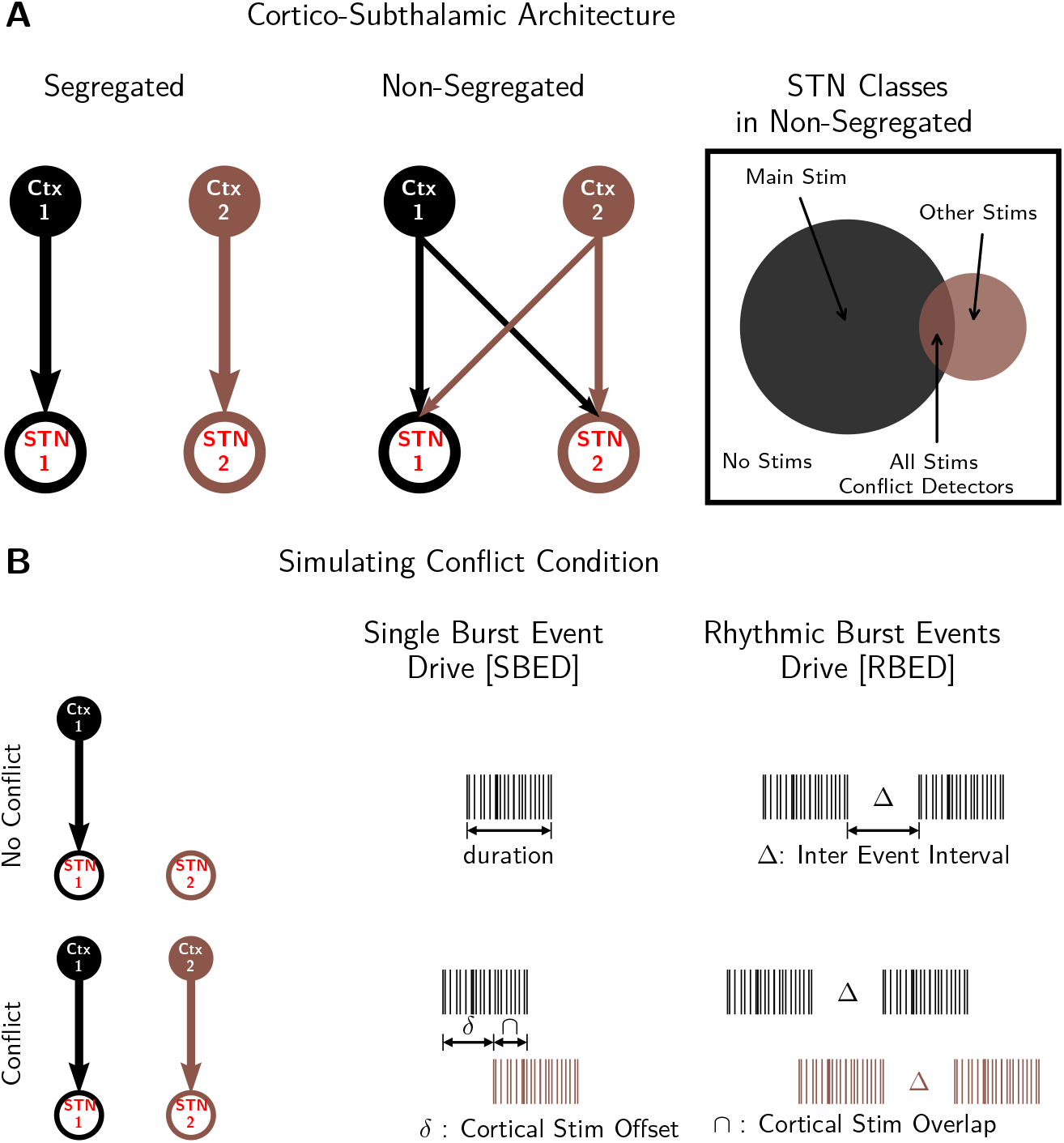
**A:** Cortico-Subthalamic network architecture. *Left:* Segregated information flow. Cortical subpopulations connect to specific STN subpoplations, precluding direct information sharing. *Middle:* Non-segregated information flow. Cortical subpolualtions connect primarily to their corresponding STN subpopulations with high probability, whilst allowing connections to other STN subpopulations, with lower probability. *Right:* Different STN classes based on stochastic cortical-STN connectivities. *Classes:* ‘MainStim’ - STN units that receive only from their own cortical subpopulation. ‘OtherStims’ - STN units that receive from only one cortical subpopulation that is not their own. ‘NoStims’ - STN units that do not receive any cortical inputs. ‘AllStims’ - STN units that receive from more than one cortical inputs, acting as *conflict detectors.* **B:** Simulating conflict condition. *Upper:* No Conflict - only one cortical subpopulation active. *Lower:* Conflict - co-activation of two cortical subpopulations, with offset *δ. Left:* Stimulation schematics. *Middle:* Single Burst Event Drive. *Right:* Rhythmic Burst Events Drive.

Given its role in theta resonance, we further examined the mechanism of this triphasic response. We analyzed individual STN membrane potential traces (Figure 4B), and found that the pause in the triphasic response reflected a depolarization block induced by the rising amplitude of the NMDA current, followed by a membrane potential recovery period aligning with the decay timecourse of the NMDA current (Figure 4C). Since the NMDA currents dominated the generation of the resonant theta frequency response, the duration of theta band resonance corresponded with the pause duration (Figure 4D). Only after NMDA currents decayed sufficiently, the GPe-mediated GABAergic currents became effective, leading to subthreshold oscillations in the membrane potential, followed by burst firing - completing the triphasic response.

To further understand the separable roles of NMDA and GABA currents in this cortically driven triphasic response, we varied both of their conductances. We first confirmed that increases in GPe GABAergic conductance increased the pause durations (Hamani 2004) and reduced the burst activities, without affecting initial spiking (data not shown). Moreover, larger NMDA conductances also prolonged the silence period. Finally, the post-silence bursts are thought to reflect rebound bursting due to T-type calcium currents activated by removal of GPe-mediated hyperpolarization (Bevan et al. 2006, 2007; Hamani 2004; Magill et al. 2006). Our investigations contribute an additional mechanistic explanation for such triphasic spiking activity. Congruent with the extant interpretations, it is initiated by excitatory AMPA currents that promote initial spiking; the onset of pause is triggered by synchronous recruitment of GABA activities from GPe. However, our model suggests that the pause is sustained beyond this inhibition due to the depolarization block induced by NMDA currents that weaken the effects of K^+^ currents, thereby preventing the hyperpolarization necessary for action potential generation. While this prediction needs to be experimentally tested, it accords with previous studies (Barraza et al. 2009; Wilson et al. 2004) that report depolarization block in the STN and their link to potassium currents. We observed that as long as cortical spiking persisted, the membrane potential was held at around −20 mV while the NMDA current accumulated. Once cortical spiking ceased, the NMDA current started to relax with a slow timecourse, prolonging the pause. Since the latter dynamics did not feature hyperpolarization, rebound bursting was due to depolarization activation of L-type calcium currents. See Beurrier et al. (2001) for alternative mechanisms underlying the silence period following high frequency stimulations, and Chiken and Nambu (2016), Hamani et al. (2017), Magariños-Ascone et al. (2002), and Perlmutter and Mink (2006) for related ‘depolarization block’ mechanisms involved in STN Deep Brain Stimulation in PD.

### Theta power is strongly modulated by conflict and cortical to STN topography

With a clearer understanding of how NMDA contributes to theta power modulation, we tested how such modulation may be additionally influenced by cortico-STN topography (see section Cortico-STN Architecture, Figure 5A), and response conflict (Figure 5B). To simulate response conflict, we included two populations of presumed cortical motor units (representing mutually incompatible responses), based on a definition of conflict as Hopfield energy (Botvinick et al. 2001; Frank 2006; Yeung et al. 2004). In the Low conflict condition, only one of these populations was active, but in a High conflict condition, they were co-active with increasing temporal overlap (Figure 5B). We varied topography of the cortical inputs, ranging from completely segregated (each cortical population projects singly to its target STN subpopulation) to fully random connections (each STN unit is equally likely to receive inputs from any cortical population; Figure 5A). Note that the number of STN units that receive inputs from more than one population increases as the topography becomes more random and less segregated. We posited that such STN units could act as “conflict detectors” because they would detect the co-activation of multiple cortical populations, and allow us to assess whether they would impact theta power in accordance with the literature (see Introduction). Cortical activity was simulated as burst drives in the two populations as would occur during action selection (Figure 5B), either as single events [SBED] or repeated “rhythmic” burst events [RBED] (the latter representing vacillating conflict within the cortex, consistent with the literature on frontal theta and conflict).

Notably, overall theta power in the network increased with lower segregation, that is, with higher number of conflict detectors (Figure 6A). An example of such high conflict with a delay between two cortical events, *δ*, of 5 ms is shown in Figure 6B, H (two-sided Welch’s T-test, p<0.05). Note that this result is not simply due to more input; indeed, we maintained the total number of inputs to the STN to be equal across all levels of segregation. Moreover, in the “no conflict” condition, theta was actually maximized by the highest segregation level, thus suggesting that topography favoring conflict detectors enhances theta only when conflict is actually present.

**Figure 6:**
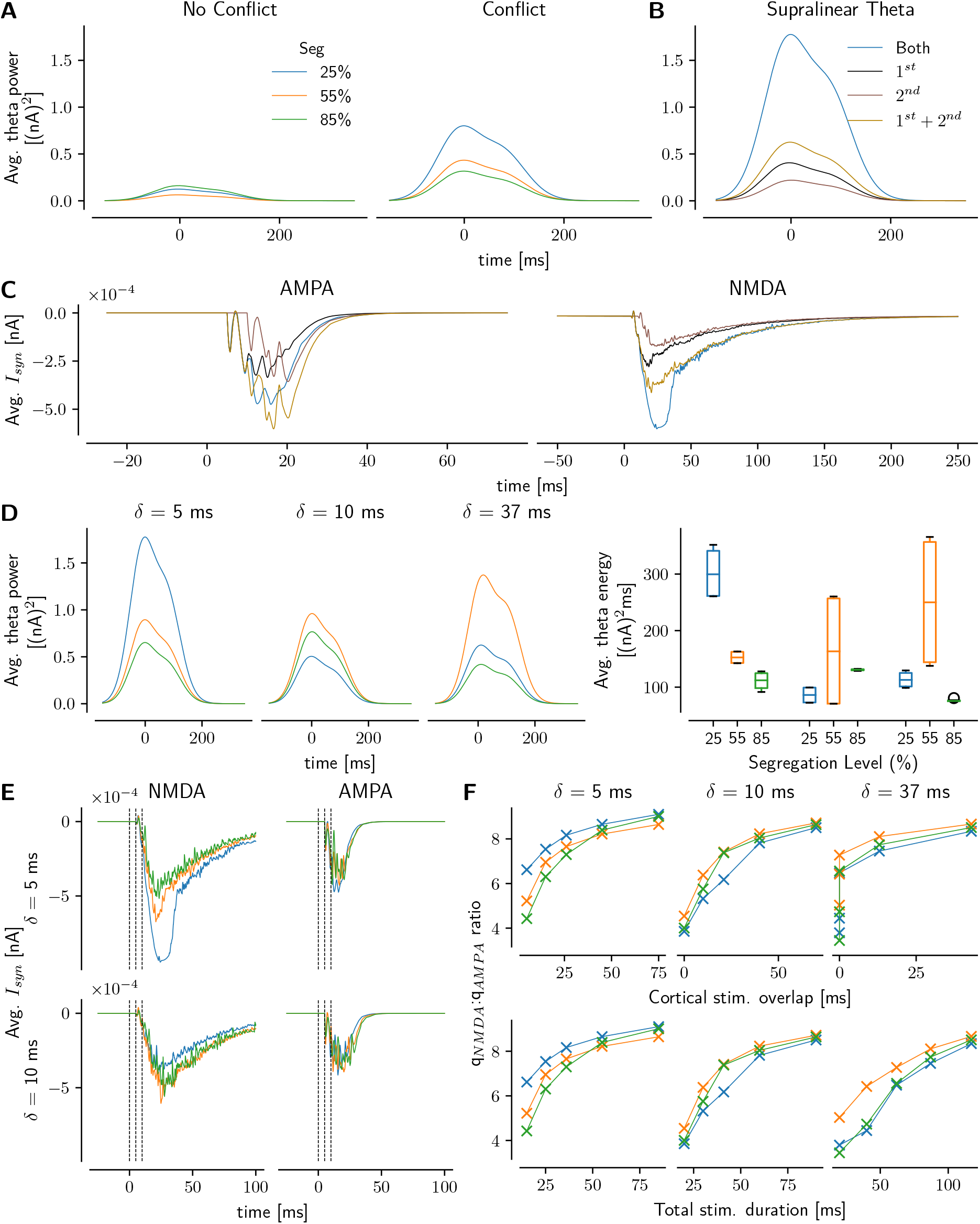
**A:** Average theta power across all STN units that received cortical inputs in *Left:* No Conflict and *Right:* Conflict conditions. **B:** Average theta power across different STN classes. Subpopulations ‘1*^st^*’ and ‘2*^nd^*’ received from either the first or the second cortical subpopulations only. ‘Both’, *STN conflict detector units*, received from both cortical drives. ‘1*^st^* + 2*^nd^*’ shows the (linear) average power from subpopulations ‘1*^st^*’ and ‘2*^nd^*’ lumped together. **C:** Average synaptic currents, *Left:* AMPA and *Right:* NMDA, across STN detector units. Color coding same as in B. **D:** Effects of cortical stimulation offset, *δ*, on average theta *Left:* power and *Right:* energy. **E:** Contribution of glutamatergic currents, *Left:* NMDA and *Right:* AMPA, towards theta power, shown for *Top: δ* = 5 ms and *Bottom: δ* = 10 ms. Broken lines represent timepoints 0, 5 and 10 ms. **F:** Nonlinear effects of cortical stimulation overlaps and durations, and segregation on the relative charge contributions of NMDA and AMPA currents observed during theta activity.

### NMDA currents enhance conflict-induced theta via supralinear summation

These results recapitulate the dominant finding in the human STN LFP literature (Cavanagh et al. 2011, 2014; Herz et al. 2016; Zavala et al. 2014; Zavala et al. 2013, 2016), whereby response conflict increases STN theta power. Our network model allows us to investigate whether such theta modulation was preferentially increased in conflict detectors. We thus computed the average theta power in separate STN subpopulations: those receiving inputs from single cortical drives 1*^st^* and 2*^nd^* and those receiving from both, 1*^st^* + 2*^nd^*. We found that theta power was indeed preferentially elevated in conflict detectors at all segregation levels. This finding stems from a supralinear summation afforded by the biophysical properties of the STN network: theta power in conflict detectors was greater than the sum of power in two populations that each received from only one cortical feed (see Figure 6B for an example).

To understand the underlying electrophysiological mechanisms for such supralinear summation, we analyzed the different glutamatergic currents, AMPA and NMDA (Figure 6C). We found that NMDA currents generated in the conflict detectors were indeed greater than the sum of the currents from the individual subpopulations receiving single cortical inputs. This pattern was not seen in AMPA currents (in fact, it was somewhat reversed, with smaller currents in the conflict detectors compared to segregated populations). This analysis suggests that response conflict increases STN theta via supralinearity mediated by NMDA currents in conflict detectors.

To further investigate how the NMDA-based dynamics of the conflict detectors interact with cortical spiking, we manipulated the durations [10-125 ms] and temporal offsets (delays) *δ*, [10-37 ms] (Figure5B) of the cortical stimulations, and consequently, the temporal overlap (i.e. conflict as defined above). We found that theta power was maximized for the lowest delays (highest overlap) in the unsegregated network (most conflict detectors). In contrast, a segregation level of 55% was more apt at theta maximization for higher delays, *δ* ≥ 10 ms. Figure 6D shows theta power and theta energy for SBED durations of 10 ms; these results are robust for SBED durations of up to 80 ms.

To understand how delay, *δ*, is critical in encoding conflict and impacting theta, and how it interacts with segregation, we analyzed the glutamatergic currents, and found that again, NMDA currents but not AMPA currents, are directly modulated by *δ*, Figure 6E. In particular, the lower the delay, the higher the opportunity for NMDA currents from the different SBEDs to integrate supralinearly, thus yielding higher theta with lowest segregation. Note that AMPA currents show no modulation with respect to either delays or segregation levels. This supports the idea that high conflict condition is encoded by the relative timing of incoming cortical signals, and mechanistically realized through the nonlinear NMDA dynamics.

It was surprising that only NMDA and not AMPA would show modulations with respect to segregation, despite being driven by identical cortical signals. Since this is due to their respective excitatory postsynaptic current rise and decay kinetics and the membrane potential synaptic conductance modulations (see section Synaptic Tuning), we characterized their differential effects by computing the ratio of overall charge transfer of NMDA to AMPA, q_NMDA_:q_AMPA_ (Figure 6F). At the lowest delay of 5 ms, the ratio increased with cortical signal overlap, ⋂ (see Figure 5B). Moreover, when overlap was low, there was a positive correlation between the ratio and number of conflict detectors (hence inverse relationship with segregation levels). However, at higher delays, *δ* ≥ 10 ms, mid-level segregation was better at reflecting the contrasting contributions.^3^

These results are notable in that the literature implicates higher theta powers being linked to higher response-time levels during response conflict, while this pattern is not seen - and indeed is often reversed - in low conflict situations (Cavanagh et al. 2011; Herz et al. 2016). We return to this issue with a plausible explanation in the discussion.

### Rhythmic Burst Events at theta frequency maximize STN spiking and theta power

Above, we observed that STN theta power could be expressed either when cortical inputs themselves oscillate in the theta range [RSSD], or when they consist of single burst events [SBED], especially with conflict. However, we also noted that higher theta power was associated with longer pauses in the triphasic spiking response. This latter result is counterintuitive, as several studies have reported *increased* STN spike rates during conflict, which correlate with larger decision thresholds (Herz et al. 2016; Isoda and Hikosaka 2008; Zaghloul et al. 2012). To investigate a possible reconciliation of these observations, we first noted that in our model, STN bursting occurred after NMDA currents decayed, which also signaled the end of theta band resonance. We then assessed the impact of prolonged response conflict in which rhythmic and burst event modes are combined [RBED], representing a situation in which conflict persists for multiple cycles in cortex (Figure 5B), consistent with the cortical literature (Bartoli et al. 2018; Zeng et al. 2021). We tested this notion by presenting repetitions of high stimulation burst events (50 ms duration, 3 ms inter burst interval, 12 burst events), where the events are repeated with different intervals Δ tested at 75, 125, 200, 350 and 450 ms. Notably, time periods from 125 through 250 ms fall in the theta range [4-8 Hz] period.

We first tested this rhythmic event [RBED] spiking protocol in the ‘no-conflict’ condition as above, and observed that, unlike in the previous case, theta power did not differ significantly across segregation levels (results not shown). Next, we presented two simultaneous rhythmic event drives to examine the influence of conflict (Figure 7A *Top*). Theta power was higher for every inter burst interval as compared to the ‘no-conflict’ condition, as expected. The instantaneous theta power was characterized by a prototypical initial peak, followed by an oscillatory period, the amplitude of which increased with IEI (Figure 7A). As previously, theta amplitudes were strongest for the lowest segregation levels, across all IEI, Δ. During the oscillatory instantaneous theta period, we observed sustained theta amplitude levels that peaked when Δ was in theta range time period [125, 250 ms]. When Δ was higher than that, instantaneous theta would rise but then decay to 0, leading to lower overall sustained power.

**Figure 7:**
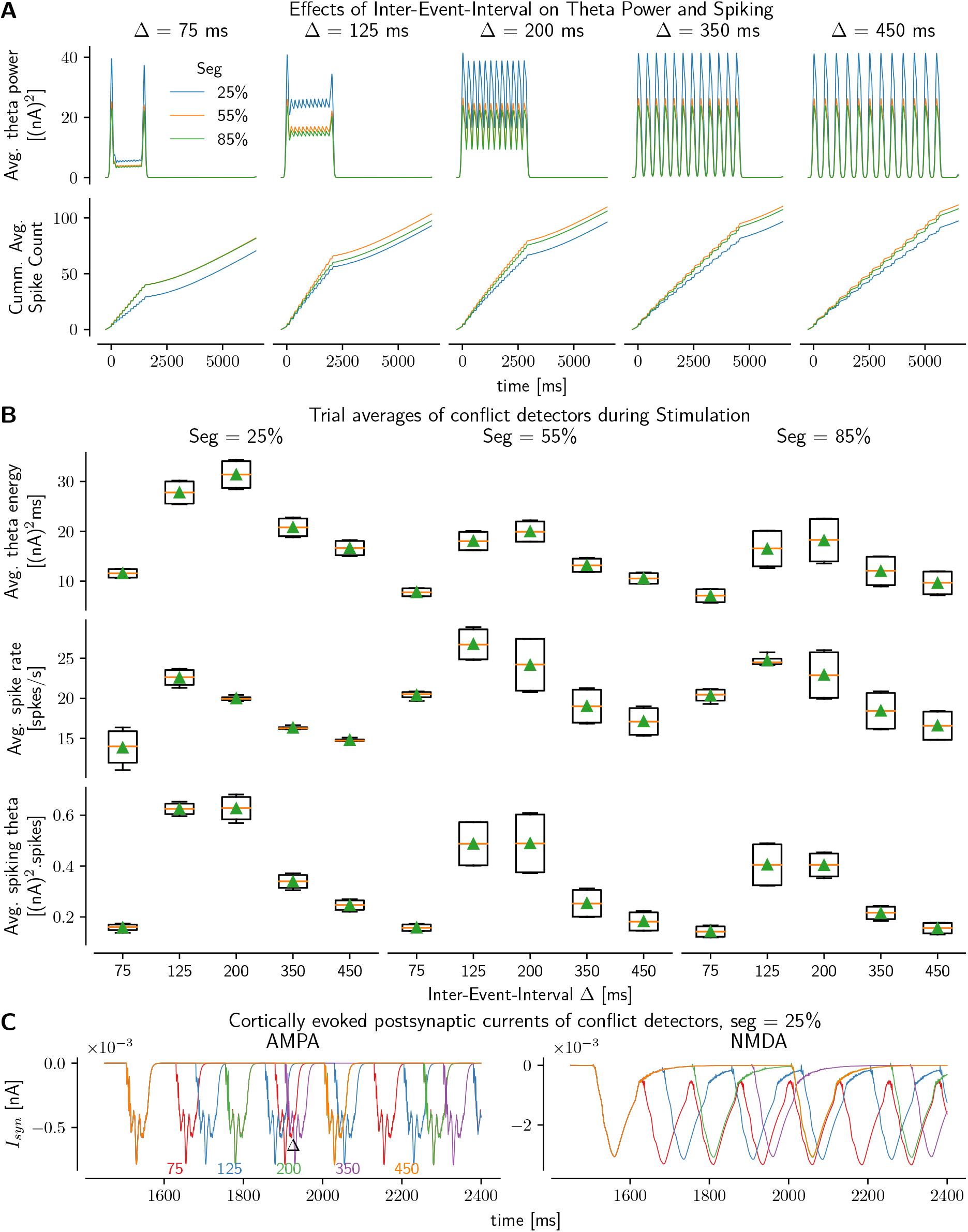
Effects of Inter-Event-Intervals, Δ, in Rhythmic Burst Events Drive cortical stimulations and segregation on STN conflict detector units. **A:** *Top:* Average theta power. *Bottom:* Cummulative average spike counts. **B:** Trial averages of conflict detectors during RBED stimulation period. *Top:* Average theta energy. *Middle:* Average spiking. *Bottom:* Average theta spiking (theta × spiking). Within each segregation level, effects of Δ on theta energy, spiking and theta-spiking are significant, p < 0.05, between theta-IEI [125, 250] and non-theta-IEI groups [75, 350, 450]. **C:** Average cortically driven post-synaptic currents of conflict detectors, shown here for the lowest segregation level 25 %.

To better characterize these nonlinear dynamical effects, we computed the overall theta energy (trial-averaged instantaneous theta power integrated over the stimulation period). First, as noted above, theta energy was highest in the lowest segregation level across all Δ and generally decreased with higher segregation (Figure 7B *Top*), though the effect of segregation was not significant (consistent with the results in Figure 6F where we observed that longer stimulation durations would eventually blur the segregation distinctions). Second, within each segregation level, Δ in theta range [125-250 ms] produced largest overall theta energy, as compared to non-theta ranges [75, 350, 450 ms]. The results were significant (two-tail Welch’s T-test, p < 0.05) between these two range groups. These were due to the fact that the theta-range promoted more elevated sustained theta, while non-theta ranges produced lower sustained theta (Δ = 75 ms) or instantaneous theta dropping to zero (Δ = 350, 450 ms) that decreased their contribution to theta rhythmicity.

Finally, since STN spiking increases during response inhibition (Fife et al. 2017; Isoda and Hikosaka 2008; Schmidt et al. 2013) and with conflict-induced elevations in decision thresholds (Herz et al. 2016), we tested the effects of IEI on spiking. We found that average spike rates of conflict detectors during cortical stimulation were maximized for Δ lying in the theta period range [125-250 ms] across all segregation levels (Figure 7A *Bottom*, B *Middle*), with results being significant between the two range groups (two-tail Welch’s T-test, p < 0.05). While there was a clear monotonic trend for theta energy as regards segregation, we observed more variable effects on spiking among segregation levels with either 25 % or 55 % generally yielding maximum spiking. In fact, this is consistent with the above observations that spiking and theta power are not always positively correlated, the more so as NMDA conductances increase.

We thus defined a new metric, theta-spiking, by taking the product of theta energy and spiking rates (Figure 7B *Bottom*), to understand under what conditions both theta and spiking rates would be *concurrently* maximized, which would be consistent with response conflict neural dynamics. First, we observed that theta-spiking was maximized when Δ was in theta range [125-250 ms], with significant differences (two-tail Welch’s T-test, p < 0.05) between the two range groups as observed for both theta energy and spiking rates separately. Moreover, segregation showed a clear monotonic trend, with lower segregation level of 25 % maximizing theta-spiking while 85 % minimizing it.

To understand the underlying biophysical mechanisms mediating these processes, and since theta power here is a direct monotonic reflection of the underlying cortically-driven postsynaptic currents, we again focus on the net glutamatergic responses (Figure 7C). Notably, for NMDA kinetics, there was a non-monotonic relationship with Δ, whereby its dynamics favor IEI lying in theta range [125-250 ms]. IEIs lower than that lead to aliasing of cortical NMDA currents, whereas those larger than the theta range exhibit no aliasing but have extended periods of silence. Δ = 200 ms lies in an optimal range that not only prevents aliasing, which enhances theta power, but also allows the NMDA currents to decay enough to basal levels to eventually promote higher spiking rates. In contrast, AMPAergic response to burst spiking remained invariant with IEI, both in trajectory and durations (about 80 ms), owing to their fast rise and decay kinetics in conjuction with the voltage-dependent synaptic conductance modulation (see Figure 1G, section Synaptic Tuning).

In sum, we observe that high conflict, high overlap and the presence of conflict detectors maximize theta. NMDA dynamics provide a filtering effect of the stochastic network spiking, thus robustly generating theta in these cases. In contrast, STN spiking is more susceptible to ongoing network stochastics, occurring mainly when NMDA currents are relatively low at the cellular level, but nevertheless maximized when cortical inputs are in the theta range. These results are consistent with observations that cortical theta is Granger causal to STN theta (Zavala et al. 2014), while also accounting for STN-GPe cellular and network dynamics that mediate changes in spiking.

## Discussion

We provide a novel theoretical framework for mechanistically interpreting the growing body of evidence linking STN theta oscillations to conflict and decision threshold adjustment (Cavanagh et al. 2011; Frank et al. 2015; Ghahremani et al. 2018; Green et al. 2013; Herz et al. 2017; Herz et al. 2016; Kelley et al. 2018; Wessel et al. 2019; Zavala et al. 2014; Zavala et al. 2013, 2016; Zavala et al. 2017). Such findings accord with existing neural network models of Basal-Ganglia, in which increased STN spike rates induce a transient increase in decision threshold during response conflict, by modulating thalamocortical activity (Frank 2006; Ratcliff and Frank 2012; Wiecki and Frank 2013); see also Bogacz and Gurney (2007). Our biophysical explorations of STN-GPe circuitry lend insight into the underlying mechanisms, leading to the following novel and testable predictions.

First, our model suggests that STN theta power does not spontaneously emerge, but rather requires cortical input and is strongly dependent on NMDA currents. The dynamics of cortical inputs were also critical: STN units exhibited theta band resonance if the cortical inputs themselves oscillate in theta (Cavanagh et al. 2014; Wessel et al. 2019; Zavala et al. 2014; Zavala et al. 2013), but theta also robustly emerged in response to cortical burst-events at a broad range of intensities and durations, particularly when such events overlapped (mimicking conflict). Moreover, both theta power and STN spiking were elevated and prolonged during rhythmic burst-events, representing a state of conflict in which cortical motor plans vacillate in the theta range. These results accord with findings that individual STN neurons exhibit increased spike rates to conflicting evidence (Isoda and Hikosaka 2008; Zaghloul et al. 2012), and that cortical theta is Granger causal to STN theta (Zavala et al. 2016). Finally, theta band resonance concomitant with spiking was also strongly modulated by architectural constraints, with maximal response when cortical inputs provided divergent connectivity to multiple STN subpopulations. Analysis of the underlying mechanisms of such effects revealed an NMDA-dependent supralinear response in STN “conflict detector” units.

### NMDA Mechanisms

The slow time course of NMDA currents, biophysically constrained from the literature, was responsible for the modulation of theta. Theta band resonance emerged even in response to single burst-events, and magnified during rhythmic bursts, due to the time period of the envelope of the decaying phase of the NMDA current. Note that we did not tune any parameters to reproduce these experimental observations; they emerged from the biophysical constraints. Notably, while theta band resonance is often described in terms of low frequency “oscillations”, our investigations align with biophysical models demonstrating how resonance can also emerge from burst-events and not just rhythmic activities (Jones 2016).

On the applied end, such findings imply that it may be possible to enhance STN function in tasks that require cognitive control by targeting local NMDA receptors, particularly the NR2D subtype, which are most prevalent in STN and GPe (Callahan et al. 2020; Yi et al. 2020). Indeed, these authors have recently developed a positive allosteric modulator of NR2D, and showed that it effectively reduced premature responding in a conflict task in rats - precisely the phenotype that is elevated with STN lesions (Baunez and Robbins 1997; Baunez et al. 2001) and which motivated the theory that STN implements a “hold your horses” mechanism (Frank 2006). Our biophysical model provides a rationale for why NMDA currents would influence such function, and also suggests that yet stronger doses could reverse the effect (due to depolarization block). Thus, in principle, different doses of such an agent could be useful for both disorders of impulsivity (too low decision threshold) and compulsivity (too high).

### Cortico-STN communication

We found that STN theta was particularly elevated when multiple cortical populations targeted individual STN units and overlapped in time (i.e. conflict). The primate STN is topographically organized by functional domains with broadly defined boundaries, but there also exist convergence zones (Alkemade et al. 2015; Haynes and Haber 2013; Kita et al. 2014; Nambu 2011). While our findings implicate STN units as conflict detectors, STN also receives direct projections from dorsomedial frontal areas (preSMA and anterior cingulate), which themselves resonate in theta during conflict (Aron et al. 2007; Frank et al. 2015; Zavala et al. 2014). Future work should explore separable roles of these cortical inputs, but our rhythmic burst simulations suggest that such cortical theta inputs may amplify STN theta and spiking. These interactions may also be related to observations that high frequency stimulation in PD - through STN excitation or silencing - can be effective by disrupting low frequency signals but preserving high frequency ones (Garcia et al. 2005).

### Reconciling the theta-response time conundrum

While many studies have replicated the finding that low frequency oscillations are related to increases in response-time and decision threshold, it is noteworthy that all of these findings were specific to high conflict task conditions (Cavanagh et al. 2011; Frank et al. 2015; Green et al. 2013; Herz et al. 2017; Herz et al. 2016; Kelley et al. 2018; Wessel et al. 2019; Zavala et al. 2014; Zavala et al. 2013, 2016; Zavala et al. 2017). If frontal and STN theta band powers were simply a “read out” of experienced conflict, then one should be able to predict response-time from theta power, irrespective of task condition. However, empirical findings have shown the opposite: within low conflict conditions, larger frontal and STN theta powers are related to *reduced* response-time (Cavanagh et al. 2011; Herz et al. 2016). Nevertheless, evidence abounds across species that STN is needed for stopping action rather than facilitating it (Aron 2007; Aron et al. 2007; Baunez et al. 2001; Fife et al. 2017; Isoda and Hikosaka 2008; Jahfari et al. 2012; Jahfari et al. 2019; Wessel et al. 2019). Our model provides plausible mechanistic reconciliations of this observation.

First, we showed that theta was observed primarily in conflict detector units. In low conflict situations, there was still an overall increase in theta with increased cortical intensity (driven by NMDA currents in individual STN populations), but note that increased theta at the population level does not necessarily involve increased spiking across the STN population. Indeed, empirical data show that STN spike rates increase in conflict conditions (Isoda and Hikosaka 2008; Zaghloul et al. 2012), and moreover, disrupting STN function only impairs response inhibition during high conflict or surprising situations (Cavanagh et al. 2011; Fife et al. 2017; Frank et al. 2007; Ghahremani et al. 2018). Notably, in our model, STN theta is related to spike output in opposite directions, depending on the period of time after stimulus onset, due to the triphasic response. Early during a decision, theta rises during the silent period, and during low conflict trials, the large evidence in cortical input may be sufficient to induce fast responding. In high conflict conditions, choices will not yet have reached threshold; the following burst spiking period, which is amplified with rhythmic cortical bursts, induces an increase in effective decision threshold. This interpretation is consistent with neural network models in which STN increases and then collapses, leading to dynamic decision thresholds over the course of choice (Ratcliff and Frank 2012). Moreover, such a dynamic threshold is normative specifically in tasks that involve a mixture of low and high conflict trials (Malhotra et al. 2018). Thus, STN theta power may speed up choice during low conflict situations simply because spiking is actually reduced early during a choice process. Finally, it is also possible that basal ganglia is not involved as much in action selection when the correct action is sufficiently salient (Brown and Marsden 1998; Cockburn et al. 2014; François-Brosseau et al. 2009).

### Limitations and Future Directions

In these studies, we have provided both biophysical and architectural mechanisms associated with theta band power modulation by focusing on cortically evoked STN-GPe interactions. We have deliberately not included other parts of the BG network including the striatum, in order to focus on the mechanisms of cortically driven theta power in STN which appear to be critical for modulating response-time during conflict tasks. This does not imply that other parts of the network are irrelevant, and indeed it would be interesting to investigate the roles of the D1 and D2 striatal pathways on GPe and STN communication, given their strong implication in motivated behavior through cortical innervations that might also converge on the STN. For a more comprehensive picture, the model could be extended to include striatal effects to investigate how the triphasic spiking responses are differentially modulated by the D1 versus D2 tug-of-war, with particular emphasis on the arkypallidal GPe subpopulation.

A paucity of studies characterizing cortical spiking during response conflict has also limited our explorations to just a few potential cortical spiking profiles. We would expect the BG-thalamocortical loop to have more dynamical effects on cortico-STN communication, which we did not account for. Moreover, though synaptic activity would monotonically correlate with LFP, we could not consider spatial filtering effects because we neither simulated the oblong arrangement in the STN nor its detailed cellular geometries, which might affect the resonant frequencies.

## Footnotes

1. Since we use only single-compartment units with no modeling of the extracellular medium, precluding any access to source and sink currents, and because LFP reflects mainly synaptic activities (Magill et al. 2004), the net synaptic currents served as a useful proxy for LFP. To contrast single STN unit LFP with no synaptic activity from GPeP, we used the net capacitative currents for the panels in Figure 2 only. This was possible because the resonant frequencies agree between both net capacitative and net synaptic currents in the lower bands (<80 Hz).

2. We also refrained from using arbitrary mappings to linearly convert from current to voltage, since we did not model the extracellular medium and its resistivity.

3. Since combinations of *δ* and SBED durations would lead to conditions of zero overlaps ⋂ = 0 ms, we repeated the same analyses against the total duration time of stimulations that were provided to the units. The results showed similar trends as above, further confirming that mid-level segregation provides a wider dynamical range as a signal integrator when the delays are large.

## References

Abdi, Azzedine et al. (2015). “Prototypic and Arkypallidal Neurons in the Dopamine-Intact External Globus Pallidus”. In: J. Neurosci. 35, pp. 6667–6688.

Alkemade, Anneke, Alfons Schnitzler, and Birte U. Forstmann (2015). “Topographic organization of the human and non-human primate subthalamic nucleus”. In: Brain Struct. Funct. 220, pp. 3075–3086.

Aron, Adam R. (2007). “The Neural Basis of Inhibition in Cognitive Control”. In: Neurosci. 13.3, pp. 214–228.

Aron, Adam R. et al. (2007). “Triangulating a cognitive control network using diffusion-weighted magnetic resonance imaging (MRI) and functional MRI”. In: J. Neurosci. 27.14, pp. 3743–3752.

Barraza, David, Hitoshi Kita, and Charles J. Wilson (2009). “Slow Spike Frequency Adaptation in Neurons of the Rat Subthalamic Nucleus”. In: J. Neurophysiol. 102, pp. 3689–3697.

Bartoli, Eleonora et al. (2018). “Temporal Dynamics of Human Frontal and Cingulate Neural Activity During Conflict and Cognitive Control”. In: Cerebral Cortex 28, pp. 3842–3856.

Baufreton, Jérôme et al. (2009). “Sparse but Selective and Potent Synaptic Transmission From the Globus Pallidus to the Subthalamic Nucleus”. In: J. Neurophysiol. 102, pp. 532–545.

Baunez, Christelle, André Nieoullon, and Marianne Amalric (1995). “In a rat model of parkinsonism, lesions of the subthalamic nucleus reverse increases of reaction time but induce a dramatic premature responding deficit.” In: J. Neurosci. 15.10, pp. 6531–41.

Baunez, Christelle and Trevor W Robbins (1997). “Bilateral Lesions of the Subthalamic Nucleus Induce Multiple Deficits in an Attentional Task in Rats”. In: Eur. J. Neurosci. 9.10, pp. 2086–2099.

Baunez, Christelle et al. (2001). “Effects of STN lesions on simple vs choice reaction time tasks in the rat: preserved motor readiness, but impaired response selection”. In: European Journal of Neuroscience 13, pp. 1609–1616.

Benhamou, Liora et al. (2012). “Globus Pallidus External Segment Neuron Classification in Freely Moving Rats: A Comparison to Primates”. In: PLoS One 7, e45421.

Beurrier, Corinne et al. (1999). “Subthalamic Nucleus Neurons Switch from Single-Spike Activity to Burst-Firing Mode”. In: J. Neurosci. 19.2, pp. 599–609.

Beurrier, Corinne et al. (2001). “High-Frequency Stimulation Produces a Transient Blockade of Voltage-Gated Currents in Subthalamic Neurons”. In: Journal of Neurophysiology 85, pp. 1351–1356.

Bevan, Mark D, Jeremy F Atherton, and Jérôme Baufreton (2006). “Cellular principles underlying normal and pathological activity in the subthalamic nucleus”. In: Curr. Opin. Neurobiol. 16, pp. 621–628.

Bevan, Mark D and Charles J Wilson (1999). “Mechanisms underlying spontaneous oscillation and rhythmic firing in rat subthalamic neurons.” In: J. Neurosci. 19.17, pp. 7617–28.

Bevan, Mark D., Nicholas E. Hallworth, and Jérôme Baufreton (2007). “GABAergic control of the subthalamic nucleus”. In: Prog. Brain Res. Vol. 160, pp. 173–188.

Bogacz, Rafal and Kevin Gurney (2007). “The basal ganglia and cortex implement optimal decision making between alternative actions.” In: Neural Comput. 19, pp. 442–477.

Botvinick, Matthew M. et al. (2001). “Conflict monitoring and cognitive control.” In: Psychol. Rev. 108, pp. 624–652.

Brown, P and CD Marsden (1998). “What do the basal ganglia do?” In: Lancet 351, pp. 1801–1804.

Bugaysen, J., I. Bar-Gad, and A. Korngreen (2013). “Continuous Modulation of Action Potential Firing by a Unitary GABAergic Connection in the Globus Pallidus In Vitro”. In: J. Neurosci. 33, pp. 12805–12809.

Callahan, Patrick M. et al. (2020). “Modulating inhibitory response control through potentiation of GluN2D subunit-containing NMDA receptors”. In: Neuropharmacology 173, p. 107994.

Carnevale, Nicholas T and Michael L Hines (2006). The NEURON book. Cambridge University Press.

Carnevale, Ted et al. (2014). “The neuroscience gateway portal: high performance computing made easy”. In: BMC Neurosci. 15, P101.

Cavanagh, James F et al. (2011). “Subthalamic nucleus stimulation reverses mediofrontal influence over decision threshold”. In: Nat. Neurosci. 14, pp. 1462–1467.

Cavanagh, James F et al. (2014). “The subthalamic nucleus contributes to post-error slowing.” In: J. Cogn. Neurosci. 26, pp. 2637–2644.

Chiken, Satomi and Atsushi Nambu (2016). “Mechanism of Deep Brain Stimulation”. In: Neurosci. 22, pp. 313–322.

Chu, Hong-Yuan et al. (2015). “Heterosynaptic Regulation of External Globus Pallidus Inputs to the Subthalamic Nucleus by the Motor Cortex”. In: Neuron 85, pp. 364–376.

Cockburn, Jeffrey, Anne G.E. Collins, and Michael J. Frank (2014). “A Reinforcement Learning Mechanism Responsible for the Valuation of Free Choice”. In: Neuron 83, pp. 551–557.

Cooper, A J and I M Stanford (2000). “Electrophysiological and morphological characteristics of three subtypes of rat globus pallidus neurone in vitro”. In: J. Physiol. 527.2, pp. 291–304.

Coulthard, Elizabeth J et al. (2012). “Distinct roles of dopamine and subthalamic nucleus in learning and probabilistic decision making”. In: Brain 135.12, pp. 3721–3734.

Deister, Christopher A. et al. (2013). “Firing rate and pattern heterogeneity in the globus pallidus arise from a single neuronal population”. In: J. Neurophysiol. 109, pp. 497–506.

Eusebio, Alexandre et al. (2009). “Resonance in subthalamo-cortical circuits in Parkinson’s disease”. In: Brain 132, pp. 2139–2150.

Fan, Kai Y et al. (2012). “Proliferation of External Globus Pallidus-Subthalamic Nucleus Synapses following Degeneration of Midbrain Dopamine Neurons”. In: J. Neurosci. 32, pp. 13718–13728.

Fife, Kathryn H et al. (2017). “Causal role for the subthalamic nucleus in interrupting behavior”. In: Elife 6.

Folk, Mike et al. (2011). “An overview of the HDF5 technology suite and its applications”. In: Proceedings of the EDBT/ICDT 2011 Workshop on Array Databases. ACM, pp. 36–47.

François-Brosseau, Félix-Etienne et al. (2009). “Basal ganglia and frontal involvement in self-generated and externally-triggered finger movements in the dominant and non-dominant hand”. In: Eur. J. Neurosci. 29, pp. 1277–1286.

Frank, Michael J et al. (2007). “Hold Your Horses: Impulsivity, Deep Brain Stimulation, and Medication in Parkinsonism”. In: Science 318, pp. 1309–1312.

Frank, Michael J et al. (2015). “fMRI and EEG predictors of dynamic decision parameters during human reinforcement learning.” In: J. Neurosci. 35, pp. 485–94.

Frank, Michael J. (2006). “Hold your horses: A dynamic computational role for the subthalamic nucleus in decision making”. In: Neural Networks 19, pp. 1120–1136.

Garcia, Liliana et al. (2005). “High-frequency stimulation in Parkinson’s disease: more or less?” In: Trends in Neurosciences 28, pp. 209–216.

Ghahremani, Ayda et al. (2018). “Event-related deep brain stimulation of the subthalamic nucleus affects conflict processing”. In: Ann. Neurol. 84, pp. 515–526.

Goldberg, J. A. (2004). “Spike Synchronization in the Cortex-Basal Ganglia Networks of Parkinsonian Primates Reflects Global Dynamics of the Local Field Potentials”. In: Journal of Neuroscience 24, pp. 6003–6010.

Green, Nikos et al. (2013). “Reduction of Influence of Task Difficulty on Perceptual Decision Making by STN Deep Brain Stimulation”. In: Curr. Biol. 23, pp. 1681–1684.

Hahn, Philip J. and Cameron C. McIntyre (2010). “Modeling shifts in the rate and pattern of subthalamopallidal network activity during deep brain stimulation”. In: Journal of Computational Neuroscience 28, pp. 425–441.

Hamani, C. (2004). “The subthalamic nucleus in the context of movement disorders”. In: Brain 127, pp. 4–20.

Hamani, Clement et al. (2017). “Subthalamic Nucleus Deep Brain Stimulation: Basic Concepts and Novel Perspectives”. In: eneuro 4, ENEURO.0140-17.2017.

Hammond, Constance, Hagai Bergman, and Peter Brown (2007). “Pathological synchronization in Parkinson’s disease: networks, models and treatments”. In: Trends in Neurosciences 30, pp. 357–364.

Hardman, Craig Denis et al. (2002). “Comparison of the basal ganglia in rats, marmosets, macaques, baboons, and humans: Volume and neuronal number for the output, internal relay, and striatal modulating nuclei”. In: J. Comp. Neurol. 445.3, pp. 238–255.

Haynes, William I. A. and Suzanne N. Haber (2013). “The organization of prefrontal-subthalamic inputs in primates provides an anatomical substrate for both functional specificity and integration: implications for Basal Ganglia models and deep brain stimulation”. In: J. Neurosci. 33.11, pp. 4804–4814.

Herz, Damian M et al. (2017). “Distinct mechanisms mediate speed-accuracy adjustments in cortico-subthalamic networks”. In: Elife 6.

Herz, Damian M. M. et al. (2016). “Neural Correlates of Decision Thresholds in the Human Subthalamic Nucleus”. In: Curr. Biol. 26, pp. 916–920.

Hines, Michael L, Andrew P Davison, and Eilif Muller (2009). “NEURON and Python”. In: Front. Neuroinform. 3.

Isoda, Masaki and Okihide Hikosaka (2007). “Switching from automatic to controlled action by monkey medial frontal cortex”. In: Nat. Neurosci. 10, pp. 240–248.

Isoda, Masaki and Okihide Hikosaka (2008). “Role for Subthalamic Nucleus Neurons in Switching from Automatic to Controlled Eye Movement”. In: J. Neurosci. 28, pp. 7209–7218.

Jahfari, S. et al. (2012). “How Preparation Changes the Need for Top-Down Control of the Basal Ganglia When Inhibiting Premature Actions”. In: J. Neurosci. 32, pp. 10870–10878.

Jahfari, Sara et al. (2019). “Cross-Task Contributions of Frontobasal Ganglia Circuitry in Response Inhibition and Conflict-Induced Slowing”. In: Cereb. Cortex 29, pp. 1969–1983.

Janssen, Marcus L. F. et al. (2017). “Cortico-subthalamic inputs from the motor, limbic, and associative areas in normal and dopamine-depleted rats are not fully segregated”. In: Brain Struct. Funct. 222, pp. 2473–2485.

Jones, Stephanie R (2016). “When brain rhythms aren’t ‘rhythmic’: implication for their mechanisms and meaning”. In: Curr. Opin. Neurobiol. 40, pp. 72–80.

Jones, Stephanie R. et al. (2009). “Quantitative Analysis and Biophysically Realistic Neural Modeling of the MEG Mu Rhythm: Rhythmogenesis and Modulation of Sensory-Evoked Responses”. In: J. Neurophysiol. 102, pp. 3554–3572.

Kelley, Ryan et al. (2018). “A human prefrontal-subthalamic circuit for cognitive control”. In: Brain 141, pp. 205–216.

Kita, H (2010). “Organization of the Globus Pallidus”. In: Handb. Basal Ganglia Struct. Funct. Ed. by Gordon Shepherd and Sten Grillner. Oxford: Elsevier Inc.

Kita, H. and S.T. T Kitai (1991). “Intracellular study of rat globus pallidus neurons: membrane properties and responses to neostriatal, subthalamic and nigral stimulation”. In: Brain Res. 564.2, pp. 296–305.

Kita, Hitoshi (2007). “Globus pallidus external segment”. In: Gaba Basal Ganglia From Mol. to Syst. Ed. by Elizabeth D Abercrombie James M. Tepper and J Paul Bolam. Vol. 160. Progress in Brain Research. Elsevier, pp. 111–133.

Kita, Takako, Pavel Osten, and Hitoshi Kita (2014). “Rat subthalamic nucleus and zona incerta share extensively overlapped representations of cortical functional territories”. In: J. Comp. Neurol. 522, pp. 4043–4056.

Kitai, S.T. and J.M. Deniau (1981). “Cortical inputs to the subthalamus: intracellular analysis”. In: Brain Res. 214, pp. 411–415.

Kreiss, D S et al. (1997). “The response of subthalamic nucleus neurons to dopamine receptor stimulation in a rodent model of Parkinson’s disease.” In: J. Neurosci. 17, pp. 6807–19.

Magariños-Ascone, C et al. (2002). “High-frequency stimulation of the subthalamic nucleus silences subthalamic neurons: a possible cellular mechanism in Parkinson’s disease”. In: Neuroscience 115, pp. 1109–1117.

Magill, Peter J. et al. (2004). “Synchronous Unit Activity and Local Field Potentials Evoked in the Subthalamic Nucleus by Cortical Stimulation”. In: J. Neurophysiol. 92.2, pp. 700–714.

Magill, Peter J. et al. (2006). “Delayed synchronization of activity in cortex and subthalamic nucleus following cortical stimulation in the rat”. In: J. Physiol. 574, pp. 929–946.

Malhotra, Gaurav et al. (2018). “Time-varying decision boundaries: insights from optimality analysis”. In: Psychonomic Bulletin & Review 25, pp. 971–996.

Mallet, Nicolas et al. (2012). “Dichotomous Organization of the External Globus Pallidus”. In: Neuron 74, pp. 1075–1086.

Mastro, K. J. et al. (2014). “Transgenic Mouse Lines Subdivide External Segment of the Globus Pallidus (GPe) Neurons and Reveal Distinct GPe Output Pathways”. In: J. Neurosci. 34, pp. 2087–2099.

Mathai, Abraham and Yoland Smith (2011). “The Corticostriatal and Corticosubthalamic Pathways: Two Entries, One Target. So What?” In: Front. Syst. Neurosci. 5.64.

Miguelez, Cristina et al. (2012). “Altered pallido-pallidal synaptic transmission leads to aberrant firing of globus pallidus neurons in a rat model of Parkinson’s disease”. In: J. Physiol. 590, pp. 5861–5875.

Nambu, A and R Llinas (1994). “Electrophysiology of globus pallidus neurons in vitro”. In: J. Neurophysiol. 72, pp. 1127–1139.

Nambu, Atsushi (2011). “Somatotopic Organization of the Primate Basal Ganglia”. In: Front. Neuroanat. 5.

Nambu, Atsushi, Hironobu Tokuno, and Masahiko Takada (2002). “Functional significance of the cortico-subthalamo-pallidal ‘hyperdirect’ pathway”. In: Neurosci. Res. 43.2, pp. 111–117.

Nambu, Atsushi et al. (2000). “Excitatory Cortical Inputs to Pallidal Neurons Via the Subthalamic Nucleus in the Monkey”. In: J. Neurophysiol. 84, pp. 289–300.

Oorschot, Dorothy E. (1996). “Total number of neurons in the neostriatal, pallidal, subthalamic, and substantia nigral nuclei of the rat basal ganglia: A stereological study using the cavalieri and optical disector methods”. In: J. Comp. Neurol. 366.4, pp. 580–599.

Parent, André and Lili-Naz Hazrati (1995). “Functional anatomy of the basal ganglia. II. The place of subthalamic nucleus and external pallidium in basal ganglia circuitry”. In: Brain Res. Rev. 20, pp. 128–154.

Park, Choongseok, Robert M. Worth, and Leonid L. Rubchinsky (2011). “Neural dynamics in Parkinsonian brain: The boundary between synchronized and nonsynchronized dynamics”. In: Physical Review E 83, p. 042901.

Pautrat, Arnaud et al. (2018). “Revealing a novel nociceptive network that links the subthalamic nucleus to pain processing”. In: Elife 7.

Pavlides, Alex, S. John Hogan, and Rafal Bogacz (2015). “Computational Models Describing Possible Mechanisms for Generation of Excessive Beta Oscillations in Parkinson’s Disease”. In: PLOS Computational Biology 11, e1004609.

Perlmutter, Joel S. and Jonathan W. Mink (2006). “DEEP BRAIN STIMULATION”. In: Annu. Rev. Neurosci. 29, pp. 229–257.

Ratcliff, Roger and Michael J Frank (2012). “Reinforcement-Based Decision Making in Corticostriatal Circuits: Mutual Constraints by Neurocomputational and Diffusion Models”. In: Neural Comput. 24, pp. 1186–1229.

Rubin, Jonathan E (2017). “Computational models of basal ganglia dysfunction: the dynamics is in the details”. In: Curr. Opin. Neurobiol. 46, pp. 127–135.

Rubin, Jonathan E. and David Terman (2004). “High Frequency Stimulation of the Subthalamic Nucleus Eliminates Pathological Thalamic Rhythmicity in a Computational Model”. In: J. Comput. Neurosci. 16, pp. 211–235.

Sadek, Ahmed R., Peter J. Magill, and J. Paul Bolam (2007). “A Single-Cell Analysis of Intrinsic Connectivity in the Rat Globus Pallidus”. In: J. Neurosci. 27.24, pp. 6352–6362.

Schmidt, Robert et al. (2013). “Canceling actions involves a race between basal ganglia pathways”. In: Nat. Neurosci. 16, pp. 1118–1124.

Schroll, Henning and Fred H Hamker (2013). “Computational models of basal-ganglia pathway functions: Focus on functional neuroanatomy”. In: Front. Syst. Neurosci. 7.122.

Schwab, Bettina C. et al. (2013). “Synchrony in Parkinson’s disease: importance of intrinsic properties of the external globus pallidus.” In: Front. Syst. Neurosci. 7, p. 60.

Sivagnanam, Subhashini et al. (2013). “Introducing the Neuroscience Gateway.” In: IWSG.

Terman, D et al. (2002). “Activity patterns in a model for the subthalamopallidal network of the basal ganglia.” In: J. Neurosci. 22, pp. 2963–76.

Towns, J. et al. (2014). “XSEDE: Accelerating Scientific Discovery”. In: Computing in Science & Engineering 16, pp. 62–74.

Verbruggen, Frederick and Gordon D. Logan (2008). “Automatic and controlled response inhibition: Associative learning in the go/no-go and stop-signal paradigms.” In: J. Exp. Psychol. Gen. 137.4, pp. 649–672.

Wessel, Jan R, Darcy A Waller, and Jeremy DW Greenlee (2019). “Non-selective inhibition of inappropriate motor-tendencies during response-conflict by a fronto-subthalamic mechanism”. In: Elife 8.

Wiecki, Thomas V and Michael J Frank (2013). “A computational model of inhibitory control in frontal cortex and basal ganglia.” In: Psychol. Rev. 120, pp. 329–355.

Wilson, C J (2010). “Subthalamo Pallidal circuit”. In: Handb. brain microcircuits. Ed. by Gordon Shepherd and Sten Grillner. Oxford: Oxford University Press.

Wilson, C J (2013). “Active decorrelation in the basal ganglia”. In: Neuroscience 250, pp. 467–482.

Wilson, Charles J. et al. (2004). “A Model of Reverse Spike Frequency Adaptation and Repetitive Firing of Subthalamic Nucleus Neurons”. In: J. Neurophysiol. 91, pp. 1963–1980.

Wylie, Scott A et al. (2010). “Subthalamic nucleus stimulation influences expression and suppression of impulsive behaviour in Parkinson’s disease”. In: Brain 133.12, pp. 3611–3624.

Yeung, Nick, Matthew M. Botvinick, and Jonathan D. Cohen (2004). “The Neural Basis of Error Detection: Conflict Monitoring and the Error-Related Negativity.” In: Psychol. Rev. 111, pp. 931–959.

Yi, Feng et al. (2020). “PTC-174, a positive allosteric modulator of NMDA receptors containing GluN2C or GluN2D subunits”. In: Neuropharmacology 173, p. 107971.

Zaghloul, K. A. et al. (2012). “Neuronal Activity in the Human Subthalamic Nucleus Encodes Decision Conflict during Action Selection”. In: J. Neurosci. 32, pp. 2453–2460.

Zavala, B. A. et al. (2014). “Midline Frontal Cortex Low-Frequency Activity Drives Subthalamic Nucleus Oscillations during Conflict”. In: J. Neurosci. 34, pp. 7322–7333.

Zavala, Baltazar et al. (2013). “Subthalamic Nucleus Local Field Potential Activity during the Eriksen Flanker Task Reveals a Novel Role for Theta Phase during Conflict Monitoring”. In: J. Neurosci. 33.37, pp. 14758–14766.

Zavala, Baltazar et al. (2016). “Decisions Made with Less Evidence Involve Higher Levels of Corticosubthalamic Nucleus Theta Band Synchrony”. In: J. Cogn. Neurosci. 28, pp. 811–825.

Zavala, Baltazar A, Anthony I Jang, and Kareem A Zaghloul (2017). “Human subthalamic nucleus activity during non-motor decision making”. In: Elife 6.

Zeng, Ke et al. (2021). “Fronto-subthalamic phase synchronization and cross-frequency coupling during conflict processing”. In: NeuroImage 238, p. 118205.

